# Active microbial communities and their extrachromosomal elements link organic matter degradation to methane cycling in anoxic sediments

**DOI:** 10.64898/2026.04.10.716911

**Authors:** Bledina Dede, Hanna Zehnle, Emilie Skoog, Taylor Priest, Karin Beck, Helmut Bürgmann, Marie C. Schoelmerich

## Abstract

Anaerobic carbon transformation in freshwater sediments drives substantial methane emissions globally, yet the microbial taxa linking complex carbon degradation to methane production remain poorly characterized. Here, we combined metagenomics with the first metatranscriptomic dataset from the anoxic sediments of meromictic Lake Cadagno (Swiss Alps) to identify the active microbial clades, metabolic pathways, and extrachromosomal elements (ecDNA) across a depth gradient within the upper 56 cm of sediment. We recovered 802 species-level metagenome-assembled genomes (MAGs) spanning 66 phyla and identified a Bacteroidota clade (VadinHA17) as one of the most abundant and transcriptionally active populations in the sediment. This clade encodes and transcribes a broad range of diverse glycoside hydrolases (GH), indicating a central role in complex carbohydrate degradation. Transcriptional profiles suggest that this clade ferments organic substrates to acetate and hydrogen, which are key substrates for methanogenesis. In line with this, the acetoclastic methanogen *Methanothrix* and hydrogenotrophic *Methanoregula* were among the most abundant and transcriptionally active archaea in the same depth layers. Beyond microbial genomes, we detected 86,905 viral OTUs (vOTUs) and 2,136 plasmid OTUs (pOTUs), with free viruses and plasmids accounting for 5-10% and 0.2% of all sequencing reads, respectively. Notably, plasmids and viruses associated with Bacteroidota VadinHA17 encode and transcribe GHs that could augment host carbohydrate-degrading capacity. Together, these findings reveal new details on how methane production in anoxic lake sediments emerges from a network spanning primary fermentation, methanogenesis and ecDNA-mediated metabolisms.

## Introduction

Freshwater sediments are globally important reservoirs of organic carbon and major sources of methane (75-175 Tg per year) (1). Under anoxic conditions, complex organic polymers are hydrolyzed and fermented into short-chain fatty acids, carbon dioxide (CO_2_) and molecular hydrogen (H_2_). These intermediates are subsequently consumed by microorganisms that mediate the terminal steps of anaerobic carbon transformation, such as sulfate- or metal-reducing bacteria and archaea, and methanogenic archaea. Although these processes are well established in engineered systems or gut microbiome (2), the specific organisms and metabolic processes actively driving organic matter degradation in natural freshwater ecosystems remain less resolved, limiting our ability to mechanistically link microbial activity to methane production in these globally relevant sources of the greenhouse gases.

Meromictic lakes provide an ideal natural ecosystem to link community structure to function because they are permanently stratified, thus maintaining persistent redox zones: the oxic mixolimnion, a narrow chemocline, and an anoxic monimolimnion (3-5). The absence of seasonal mixing within such lakes leads to a stable environment in the sediments, where anaerobic microbial processes consistently dominate. Lake Cadagno (Piora Valley, Swiss Alps) is one of the best studied meromictic lakes and a model for redox-stratified aquatic ecosystems. Its compact chemocline sustains dense blooms of anoxygenic phototrophic purple and green sulfur bacteria and hosts tightly coupled sulfur, nitrogen, iron, and carbon cycles across steep gradients (3–5).

The chemocline is shaped by physical-biological feedbacks, including bioconvection mediated by the motile purple sulfur bacterium *Chromatium okenii* and viral processes that influence microbial productivity and organic matter recycling (5). In the lake sediment, the active processes remain less understood. Previous studies have primarily focused on geochemistry, laboratory experiments, and taxonomic profiling (6–9). They revealed that younger sediments (<1,000 years) are enriched in taxa typically associated with anaerobic surface sediments (Deltaproteobacteria, Acidobacteriota and Firmicutes), whereas deeper, older sediments harbor lineages common in energy-limited subsurface environments, including Bathyarchaeia, Chloroflexota, and Atribacteria. Despite these taxonomic changes, core metabolic capacities such as carbon fixation, fermentation, and sulfate reduction appear broadly conserved throughout the sedimentary record (9). However, the existing datasets provide limited resolution in the upper meter, where biogeochemical gradients are steepest (6,9), and metagenomic data are only available from two discrete depths of 2 and 40 cm, constraining resolution of fine-scale microbial changes near the sediment-water interface.

To address this gap, we focused on the upper 56 cm of the sediment of Lake Cadagno, representing approximately 222 years of sediment accumulation. Combining depth-resolved metagenomics with the first metatranscriptomic dataset from these sediments, we aimed to (i) identify the active bacterial and archaeal populations and metabolic pathways across a depth gradient, (ii) resolve links between complex carbon degradation, fermentation, methane and sulfur cycling, and (iii) characterize extrachromosomal elements (ecDNA) including plasmids and viruses/phages and their contributions to metabolic processes. Our findings underscore the central role of the Bacteroidota clade VadinHA17 in sediment carbon cycling and reveal a metabolic connection between complex polysaccharide degradation, fermentation, and methanogenesis. Furthermore, our results suggest previously underappreciated controls of sediment carbon flow, community structure and environmental adaptation, including potential top-down control of dominant microbial clades through ecDNA activity.

## Methods

### Sediment sampling at Lake Cadagno

Two 56-cm sediment cores were collected using a gravity corer and winch (EAWAG) beneath a 20.9-21 water column at Lake Cadagno (Piora Valley, Ticino, Switzerland) on 20 July 2024. A few centimeters of overlying water were retained at the top of the corer to ensure the complete capture of the sediment core. Core 1 was processed immediately after sampling on the lake platform, while Core 2 was stored at 4 °C and processed in the laboratory within a few days. Each core was divided into six horizontal sections: 0-10, 10-20, 20-30, 30-40, 40-50, and 50-56 cm. Material from each section was homogenized thoroughly, transferred to sterile Whirl-Pak bags, and stored at -80 °C until further processing.

### DNA and RNA extraction and sequencing

From each section of both cores, 5 g sediment was taken from the Whirl-Pak bags for nucleic acid extraction. DNA and RNA were co-extracted from the same sample using the Qiagen RNeasy PowerSoil Total RNA Kit combined with the RNeasy PowerSoil DNA Elution Kit, following the manufacturer’s instructions (**Table S1**).

Twelve metagenomes (six from each core) were sequenced using paired-end 150 bp reads on the Illumina NovaSeq X Plus platform at Novogene (Munich, Germany). For metatranscriptomics, extracted RNA underwent rRNA depletion and subsequent shearing into fragments of 250-300 bp. The fragments were then reverse-transcribed into cDNA. The cDNA was subject to end repair, A-tailing, adapter ligation, size selection, and PCR amplification. All six depth sections of Core 1 yielded RNA of sufficient quality for sequencing, whereas only the first two sections (0-10 cm and 10-20 cm) of Core 2 met the quality threshold. Consequently, a total of eight metatranscriptomes were sequenced using the same paired-end 150 bp approach on the Illumina NovaSeq X Plus platform at Novogene (**Table S2**), generating approximately 50 Gbp of raw sequencing data per sample.

### Raw sequence data processing and recovery of MAGs

Raw metagenome data was quality-checked using FastQC v0.12.1 (10) and trimmed using BBduk of the BBMap package v39.01 (ktrim=r, k=23, mink=11, hdist=1, trimpolyg=20, trimq=20, qtrim=rl, minlen=105, tpe, tpo) (11) . The trimmed reads then underwent another quality check with FastQC before being assembled with metaSPAdes v4.0 (k-mers: 21,33,55,77,99, and 127) (12). Scaffolds shorter than 1,000 bp were excluded from the final assemblies (**Table S3**).

Taxonomic composition of microbial communities was assessed using two assembly-independent approaches. SingleM v0.18.3 (13) was run on the raw sequences, as specified by the developers, to estimate microbial community composition. In parallel, taxonomic profiling was performed with the mOTU-profiler v4.0.4 (default parameters) (14) to obtain a species-resolved profile of microbial communities. Using both approaches allowed us to cross-validate taxonomic patterns. Metagenome-assembled genomes (MAGs) were reconstructed from the assemblies. Further details on the binning are available in Supplementary Information (SI).

For metatranscriptome processing, raw reads were trimmed and quality-checked using the same BBduk parameters and FastQC pipeline as described above for the metagenome data. Metatranscriptomic reads were then processed following the TranscriptM pipeline v0.3.1 (15). Briefly, filtered reads were aligned to the recovered MAG dataset using Bowtie2 v2.5.4 (16), and gene-level transcript abundances were quantified using featureCounts v2.0.6 (17) in paired-end mode, restricting analysis to coding sequences (CDS features) and enabling strand-specific counting.

### EcDNA identification, abundance, and activity analysis

Viruses were identified from metagenomic assemblies generated from each sediment depth interval (0-56 cm) using geNomad v1.8.1 (18). Viral contigs ≥1 kb were extracted from each assembly and merged across depths prior to quality assessment. Viral quality and completeness were evaluated using CheckV v1.0.3 (19), and contigs shorter than 3 kb or encoding fewer than three genes were removed. Proviral and viral contigs passing quality filters were dereplicated using CD-HIT-EST v4.8.1 (20) at 95% nucleotide identity with ≥85% alignment coverage to define viral operational taxonomic units (vOTUs). The resulting vOTU genome set was used for downstream analyses.

Plasmid sequences were also identified from each sediment core layer (0-56 cm) using geNomad, and plasmid hits with a minimum scaffold length of ≥1 kb were retained. Redundant plasmid sequences were clustered and dereplicated using dRep (21) at 95% ANI and coverage 50%, retaining one representative per cluster, plasmid operational taxonomic units (pOTUs).

Further details on vOTUs and pOTUs abundance, transcription and host-linkage is given in SI.

### Community-level analysis: Estimation of abundance and transcription of genes, MAGs, and viral contigs

The gene catalogue was created (details in SI) and profiled across all metagenomes and metatranscriptomes by aligning reads with bwa_mem v0.7.19 (22). Alignments were filtered to retain reads with a >=95% sequence identity and >=80% coverage. To ensure robust identification and downstream abundance estimations, only genes with >=95% horizontal coverage of reads in a sample were retained. The vertical coverage of genes was then defined as the median read depth per base. For metagenomes, the number of genomes sequenced was estimated as the median of the summed vertical coverage of 40 universal single-copy marker-genes per sample. For metatranscriptomes, the same approach was used to estimate the number of transcribed genomes in each sample. The vertical coverage of each gene cluster was then normalized by the number of genomes detected to obtain an estimate of copies per genome.

Gene expression was inferred by normalizing the metatranscriptomic copies per genome by the corresponding metagenomic copies per genome, followed by log_2_ transformation. This metric represents transcription relative to gene copy number: a value of 0 indicates equal coverage in metagenomes and metatranscriptomes, while positive values indicate higher-than-expected transcription - a proxy for upregulation. Coverage and expression of protein families was estimated by aggregating gene-cluster-level information based on functional annotations. Copies per genome for each protein family were calculated as the sum of the corresponding gene clusters, and expression values were derived using the same normalization procedure described above. MAG abundance and expression were estimated by normalizing the median copies per genome across its single-copy marker-genes by the estimated number of genomes sequenced (total and transcribed), yielding the fraction of the community represented by that MAG in metagenomes and metatranscriptomes.

## Results and discussion

### Microbial community composition is dominated by Bacteroidota

Marker-gene-based profiling (mOTUs) of two 56 cm sediment cores sampled at 10 cm intervals revealed that bacterial communities were dominated by Bacteroidota (previously Bacteroidetes), Bipolaricaulota, and Desulfobacterota, whereas archaeal communities were primarily composed of Halobacteriota, Thermoplasmatota, and Thermoproteota (**Figure 1A**). Bacteroidota was the most abundant and transcriptionally active phylum in the upper 30 cm of sediment, reaching up to 30% of the total community and accounting for up to 57% of transcribed genomes. The major contributor of the Bacteroidota signal was *Chlorobium*, a genus of well-characterized anoxygenic phototrophs. Their high transcription in the lake sediment was particularly noteworthy, given that these organisms are typically associated with the sunlit layers of the water column. Below 20 cm depth, the community shifted toward a greater representation of archaeal lineages. Thermoproteota increased in relative abundance (up to 2%) and showed particularly high transcriptional activity (up to 12%) in deeper layers. A substantial fraction of the community in deeper layers was unclassifiable with mOTUs, indicating that many taxa in these layers represent species which are missing from the reference database.

**Figure 1.**
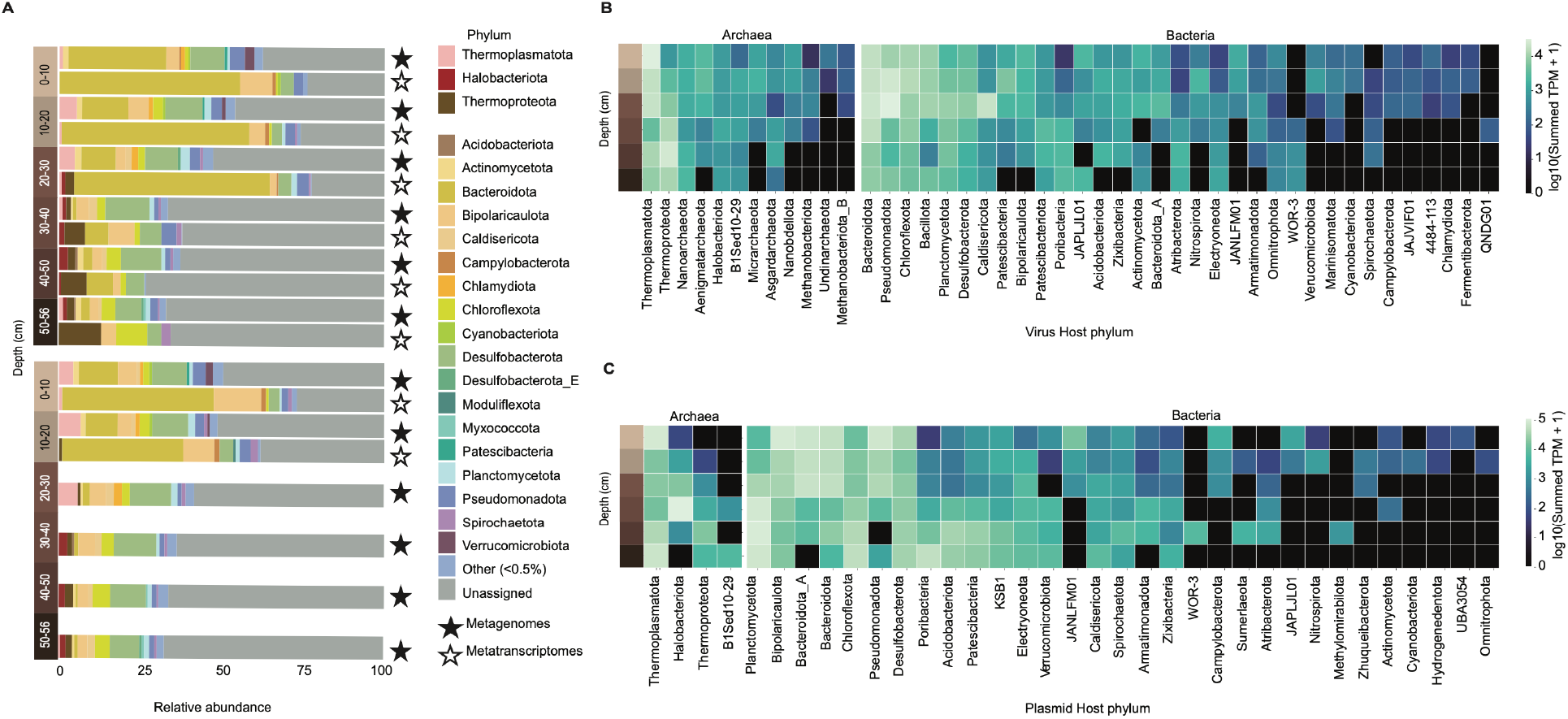
Marker-gene-based profiles of microbial community and ecDNA composition and transcriptional representation in Lake Cadagno sediments. **A.** Phylum-level taxonomic composition of the total microbial community inferred from metagenomic and metatranscriptomic datasets, grouped at a 0.5% relative abundance threshold. Taxonomic profiles were generated using mOTUs marker-gene-based profiling, and relative abundances are shown separately for metagenomes and metatranscriptomes. **B**. Transcription of viruses with predicted host associations across sediment depth. Only viruses linked to hosts were considered (7% of viruses). **C**. Transcription of plasmids with predicted host associations across sediment depth. Only plasmids linked to hosts using the iPHoP tool were considered (30% of plasmids).

An alternative taxonomic profiling approach based on 59 protein-coding marker-genes (SingleM) confirmed the dominance of Bacteroidota, Thermoplasmatota, and Bipolaricaulota at both the genomic and transcriptomic levels (**Figure S1**). Discrepancies between mOTUs and SingleM tools have been discussed in SI. Notably, SingleM profiling recovered a striking signal from Patescibacteriota (CPR) and Nanoarchaeota (DPANN), which together comprised 5.6–16.9% and 4.3–8.0% of the community, respectively, across all samples and displayed remarkable taxonomic diversity. Despite this high abundance and diversity, both groups contributed little to the metatranscriptomic signal.

Our findings are broadly consistent with previous depth-resolved investigations of Lake Cadagno sediments, which reported *Chlorobium* abundance down to 20 cm depth, while Bacteroidota exhibited elevated abundance in surface layers (6,9). Notably, *Chlorobium* was previously identified as the sole taxon encoding the sulfur oxidation genes *soxA* and *soxB* in surface sediments (9), highlighting its specialized role in sulfur-based metabolisms (23). However, these earlier studies relied only on 16S rRNA gene surveys and metagenomic data (6,9). By incorporating metatranscriptomic analyses, our study reveals that Bacteroidota, including the previously considered obligate phototrophic *Chlorobium* (24), are not only highly abundant, but also active in the absence of light.

### Diverse plasmids and viruses are actively transcribed throughout the sediment profile

Across all depths, we recovered 86,905 dereplicated, quality-filtered vOTUs and 2,136 pOTUs. While proviruses contributed ≤0.01% of total DNA reads at each depth, putatively free viral sequences accounted for ∼5-10% (**Figure S2**). In contrast, viral transcripts comprised 0.24-0.59% of transcripts and generally decreased with depth (**Figure S2**). The viral community was largely dominated by Unclassified viruses and Unclassified Caudoviricetes, several giant virus lineages, including Phycodnaviridae, Mimiviridae, and Unclassified Megaviricetes (**Figure S3**) consistent with reports from permanently stratified Lake Pavin and proglacial lake sediments from the Tibetan Plateau (25,26).

Leveraging iPHoP, we predicted putative hosts for 30% of plasmids (n = 646) and 7.1% of viruses (n = 6,210). We focused on these host-linked ecDNA for downstream interpretation. The most transcribed viruses were associated with hosts belonging to Thermoplasmatota, Bacteroidota, Pseudomonadota, and Chloroflexota (**Figure 1B**). Archaeal viruses predicted to infect Thermoplasmatota were highly expressed in the upper sediment interval (0–30 cm), whereas Thermoproteota-associated (Bathyarchaeota) viruses showed higher expression in deeper layers, highlighting a niche partitioning of their host (**Figure 1B**). This is in line with previous studies reporting Bathyarchaeota as highly abundant in the deeper layers of Lake Cadagno sediment (9). Most bacterial viruses were primarily active in surface sediments, including those linked to Bacteroidota, Pseudomonadota, and Chloroflexota (**Figure 1B**).

Notably, Patescibacteria-infecting viruses exhibited particularly high transcriptional levels in deeper layers, potentially explaining the Patescibacteria abundance (**Figure S1**), despite their comparatively low expression. In contrast, Bacteroidota exhibited high transcription of both host and viral genes. In surface sediments, the most transcribed Bacteroidota-associated viruses were affiliated with the SM23-62, *Bacteroidaceae*, and *Chlorobiaceae* clades (**Figure S4**), whereas Bacteroidota VadinHA17-associated viruses were more transcribed in deeper sediments (30–50 cm). This data indicates tight host-virus coupling, however, given the lack of temporal resolution, we cannot assess whether this reflects balanced infection dynamics or a transient phase of host decline.

Plasmid-associated sequences consistently represented ∼0.1% of total DNA reads at each depth and contributed between ∼0.01-0.10% of total transcripts, and decreased with depth, particularly below 40 cm (**Figure S2**). Plasmids were transcribed throughout all layers, with Thermoplasmatota and Thermoproteota plasmids showing the highest transcription in the deeper layers (40–56 cm). In contrast, bacterial plasmids were primarily expressed in the upper layers (notably those of Planctomycetota, Bipolaricaulota, Pseudomonadota and Bacteroidota), whereas Desulfobacteriota and Chloroflexota plasmids were mostly expressed in the deeper layers (**Figure 1C**).

Altogether, these results reveal a vast and depth-structured diversity of viruses and plasmids, pointing to a complex network of ecDNA interactions shaping the microbial community throughout the sediment profile.

### Genome-resolved metagenomics reveals high microbial diversity, with Bacteroidota among the most abundant and transcriptionally active

Metagenomic binning yielded 1,730 high-quality MAGs, which were dereplicated at the species level using a 95% ANI threshold. The resulting 802 species-representative MAGs (**Table S4**) encompassed 66 phyla, highlighting the extensive phylogenetic diversity present in these sediments, and collectively represented up to 50% of the communities. Among these, bacterial MAGs (n=603) spanned 55 phyla and archaeal MAGs (n = 199) encompassed 11 phyla (**Figure 2A, Table S5**). Consistent with the marker-gene-based community profiles (**Figure 1A**), Bacteroidota MAGs (n=57) were among the most abundant and transcriptionally active, comprising up to 4% of the community and 17% of all transcripts (**Figure 2**).

**Figure 2.**
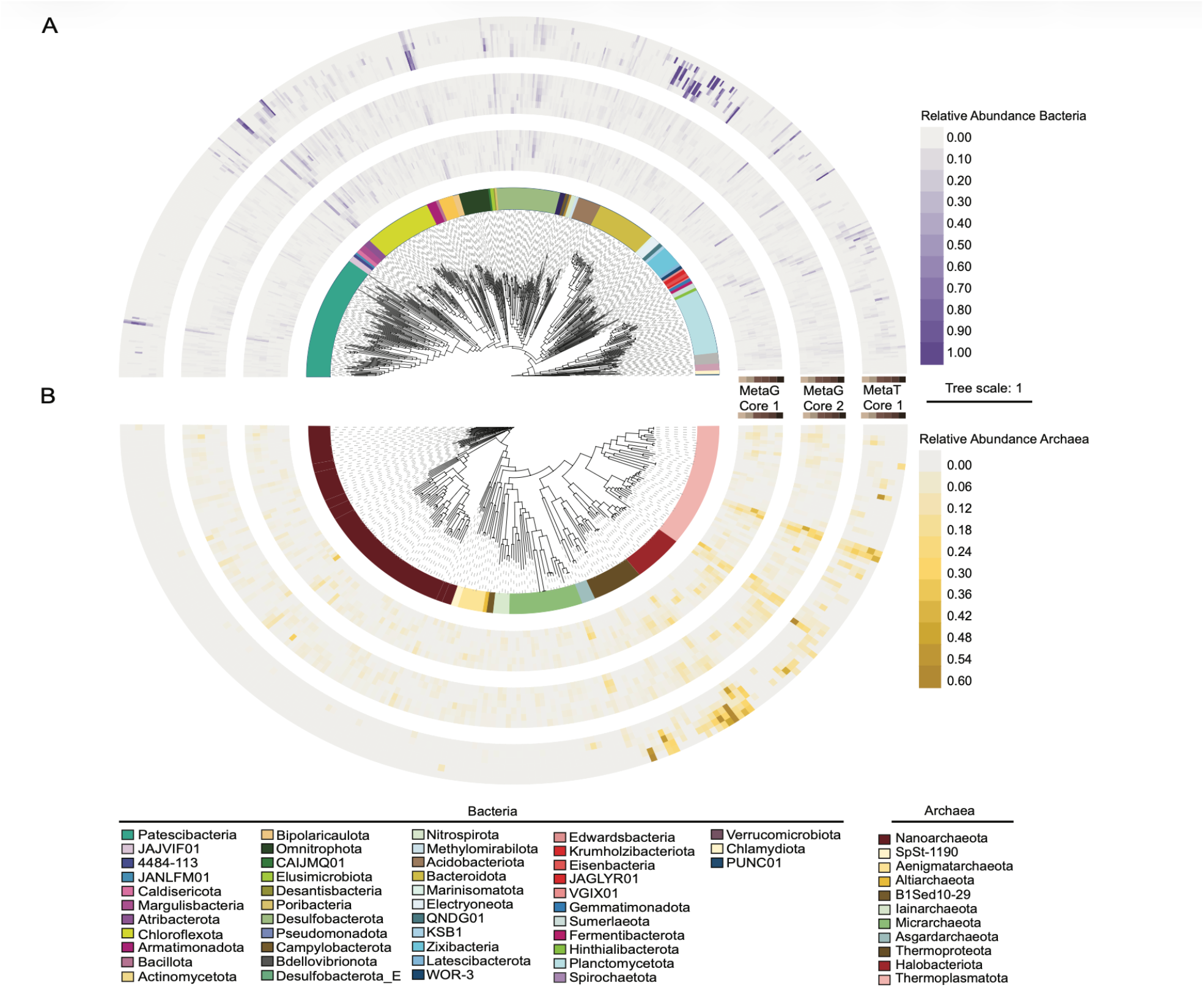
Phylogenetic trees of A. bacterial and B. archaeal MAGs. The inner layer of each tree depicts the relative abundance of MAGs in metagenomes of Core 1 and 2, while the outer ring shows the abundance in metatranscriptomes of Core 1.

Among archaea, the dataset was dominated by Nanoarchaeota (n=80), Thermoplasmatota (n=42), Microarchaeota (n=23), and Halobacteriota (n=16) (**Figure 2B**). This taxonomic composition is broadly consistent with other anoxic sediments, where archaeal communities are typically structured by a combination of symbiotic lineages, heterotrophic, and methanogenic taxa (27,28). Halobacteriota, including members of the methanogenic genera *Methanoregula* (class *Methanomicrobia*) and *Methanothrix* (class *Methanosarcinia*), increased in relative abundance with depth (**Table S6**). This vertical distribution is consistent with progressively reduced conditions favorable for methanogenesis and aligns with several studies reporting depth-stratified methanogenic communities in anoxic sediments (29,30).

Overall, bacterial and archaeal community composition shifted with depth from diverse, predominantly heterotrophic taxa in the upper sediment layers towards communities with greater representation of anaerobic archaea, including methanogens, in deeper layers.

### Heterotrophic carbon degradation dominates the metabolic landscape

To identify the most active metabolic pathways, we examined the ten most highly expressed MAGs from each depth layer (36 unique MAGs in total). These spanned diverse phyla, with Bacteroidota, primarily VadinHA17 and *Chlorobiaceae* being the most represented (**Figure 3**).

**Figure 3.**
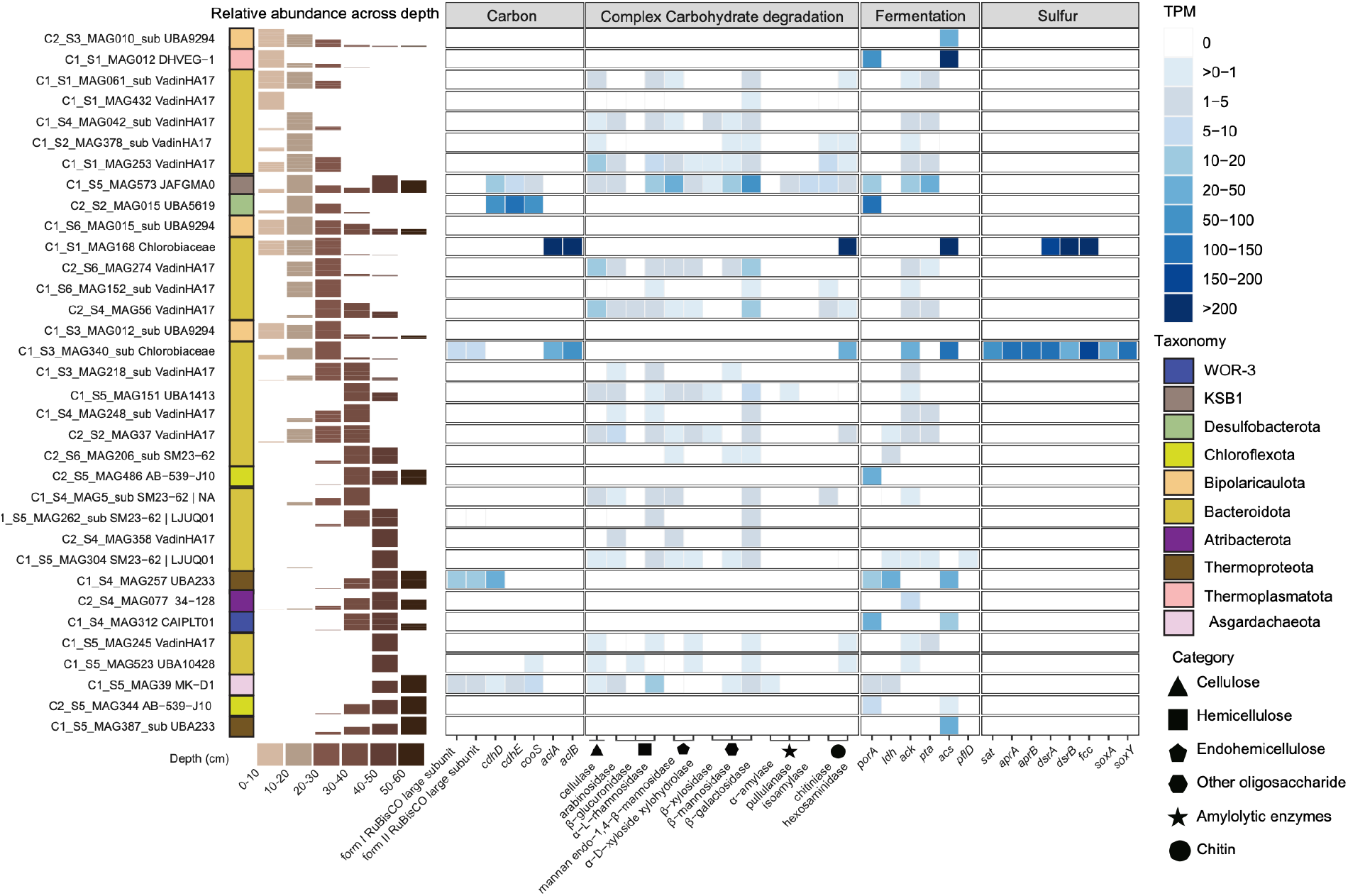
Transcriptional activity of selected functional genes across most active MAGs. For each depth interval, the ten MAGs with the highest metatranscriptomic read coverage were selected, and the colored bar on the left indicates the depth intervals in which each MAG ranked among the top ten. Gene abbreviations are as follows: RuBisCO large subunits form I and II; *cdhDE* – CO dehydrogenase/acetyl-CoA synthase; *cooS* – monofunctional anaerobic carbon-monoxide dehydrogenase; *aclAB* – ATP citrate lyase subunits A and B; *porA -* pyruvate ferredoxin oxidoreductase alpha subunit; *ldh -* L-lactate dehydrogenase; *ack* – acetate kinase; *pta* – phosphotransacetylase; *acs* – acetyl-CoA synthetase; *pflD* – pyruvate formate-lyase; *sat* – sulfate adenylyltransferase; *aprAB* – adenylylsulfate reductase subunit A and B; *dsrAB* – dissimilatory sulfite reductase subunit A and B; *fcc* – flavocytochrome C sulfide dehydrogenase; *soxAY* – sulfur oxidation pathway genes.

*Chlorobiaceae* MAGs (**Figure S6**) were notable for their metabolic versatility. All of the following pathways were actively transcribed in C1_S2_MAG168 and C1_S3_MAG340_sub (**Figure 3**): sulfide or thiosulfate oxidation via the SOX pathway, producing intra- or extracellular sulfur globules that may subsequently be further oxidized via the reverse dissimilatory sulfate reduction system (31), the membrane-bound [NiFe] hydrogenase (subgroup 1e) for hydrogen oxidation, nitrogen fixation (*nifH*), and carbon fixation via the reverse TCA cycles (*aclAB*). In addition, C1_S2_MAG168 encodes *norC*, a subunit of nitric oxide reductase, indicating a capacity for nitric oxide reduction as a respiratory electron sink in the absence of light (**Table S7**). Given this metabolic potential, *Chlorobium* could fix CO_2_ by oxidizing sulfide or hydrogen and either reducing nitric oxide or transferring electrons to a metabolic partner. This proposed strategy is supported by the fact that dark carbon fixation has been previously observed in *Chlorobium* populations within the Lake Cadagno water column (32,33). Altogether, this large repertoire suggests that *Chlorobium* in this system functions as a flexible chemolithoautotroph, capable of adapting to fluctuating redox conditions rather than being restricted to a strictly phototrophic lifestyle.

Chemolithoautotrophic pathways were also present in a few other MAGs. One Desulfobacteriota C2_S2_MAG015 and one KSB1 C1_S5_MAG573 encoded and transcribed the Wood-Ljungdahl pathway key genes (*cdhD, cdhE* and *cooS*), suggesting capacity for carbon fixation. However, although autotrophic processes are present, the most transcriptionally active community members are the heterotrophs driving carbon degradation (**Figure 3, Figure S5**).

The majority of the most active MAGs encoded a broad repertoire of CAZymes, including glycoside hydrolases (GH) and polysaccharide lyases (PL), indicating that the enzymatic breakdown of complex polysaccharides and other organic polymers is a dominant metabolic process in these sediments (**Figure S7**). Copy numbers of CAZymes reached several hundred per MAG, particularly within the Bacteroidota_VadinHA17 (**Figure S8**). For comparison, gut-associated Bacteroidota can encode 0-400 GH per genome (34), while marine sediment microbial communities typically harbor up to 16 GHs per Mbp (35), placing the VadinHA17 genomes in our lake sediments within the upper end of this range (as many as 54 GHs per Mbp). This extraordinary enzymatic repertoire highlights their capacity for polysaccharide degradation and positions them as central players in organic matter turnover. The sustained activity of these pathways is likely supported by the high seasonal rates of primary production and episodic terrestrial organic carbon input (8).

### Glycoside hydrolases are upregulated in deeper sediment layers

Given the observed dominance of heterotrophic carbon degradation processes and the high transcriptional activity of CAZymes among the most active MAGs, we expanded our analysis to investigate carbohydrate degradation across the broader community along the sediment depth gradient. To this end, we predicted CAZyme genes across all metagenomic contigs and quantified their expression by normalizing metatranscriptomic to metagenomic coverage (see Methods). This approach enabled us to resolve the taxonomic distribution and depth-dependent expression patterns of GH families, and to contextualize MAG-derived signals with contributions from the unbinned fraction of the community.

Across all depth layers, the highest CAZyme expression was observed in a subset of taxa, particularly members of the Bacteroidota and Planctomycetota (Figure 4A). While some taxa exhibited depth-specific contributions, appearing among the most highly expressed GH families only in either surface or deeper layers, these typically showed high expression in a limited number of GH families. In contrast, Bacteroidota and Planctomycetota consistently exhibited high expressions of diverse GH families across depths.

**Figure 4.**
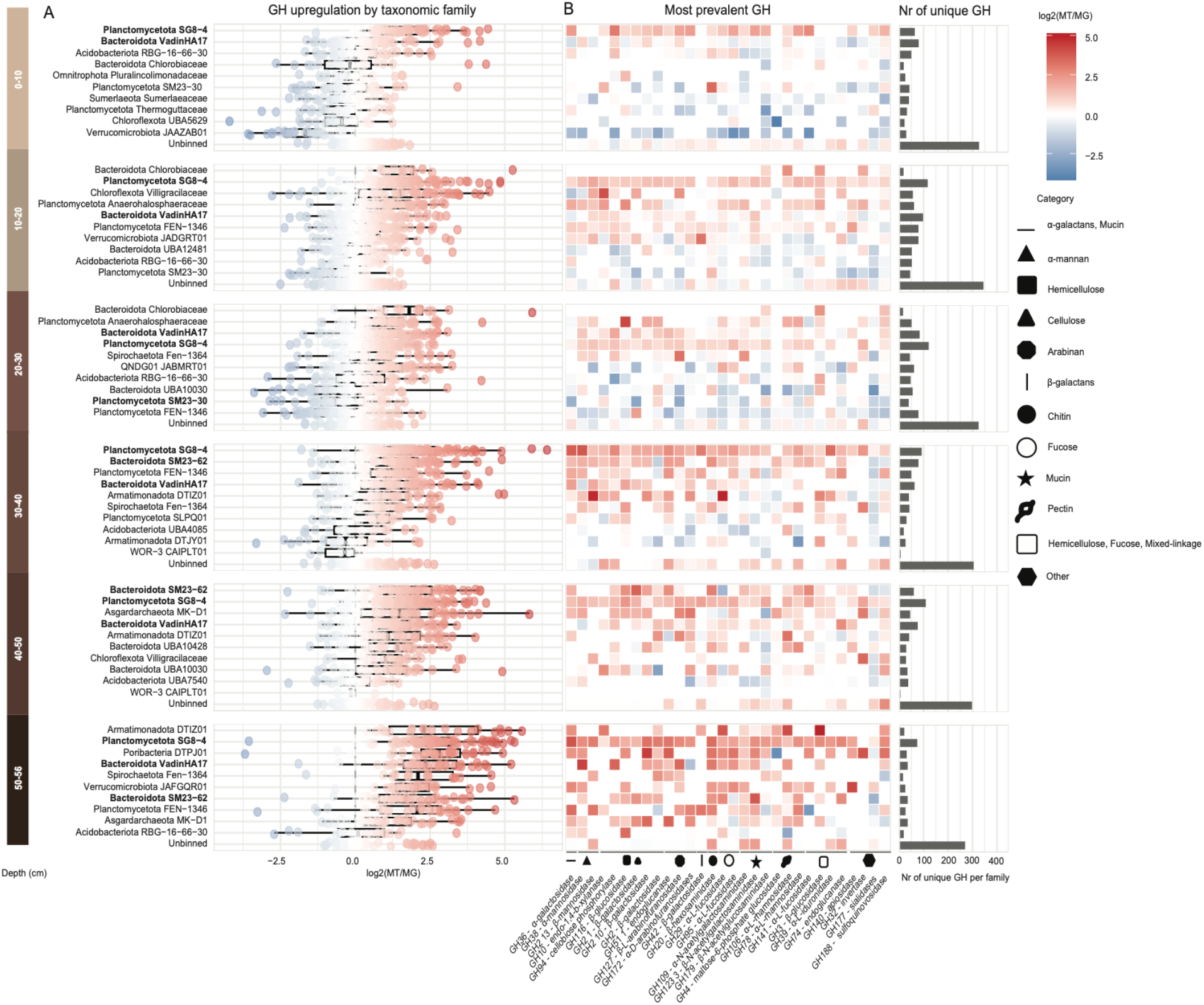
Transcriptional upregulation of glycoside hydrolase genes in Lake Cadagno sediments. **A.** Transcriptional upregulation of GH genes by taxonomic families (top 10 + unbinned). GH genes not assigned to any MAG were pooled as “unbinned GH”. **B**. The upregulation of 30 most prevalent GH families is depicted in the heatmap and the number (nr) of unique GH per family is indicated on the right hand side. Bold labels indicate Bacteroidota (VadinHA17, SM23-62) and Planctomycetota (SG8-4) families present across all layers.

Notably, we observed that both the diversity and magnitude of CAZyme transcription within these dominant Bacteroidota and Planctomycetota increased with depth, such that the majority of their CAZyme repertoires were upregulated in the deepest sediment layers (**Figure 4A**). Within Bacteroidota, GH expression was dominated by members of the SM23-62 and VadinHA17 clade, which exhibited particularly high upregulation of GH30 (α-mannosidase), GH109 (endo-acting α-N-acetylgalactosaminidase), GH20 (exo-acting β-N-acetylglucosaminidases), and GH29 (exo-acting α-fucosidases). These enzyme families target structurally complex polysaccharides typically associated with plant material, microbial biomass, and extracellular polymeric substances. Consistent with this, we also identified 14 PULs (Polysaccharide Utilization Loci) within VadinHA17-affiliated MAGs (**Table S8**), including loci predicted to target the degradation of terrestrial-derived polysaccharides. These patterns are consistent with an adaptation to complex, recalcitrant carbohydrate utilization in energy-limited, anoxic environments, in line with the previously reported enrichment of degradation-resistant carbohydrates, such as lignin, below 20 cm depth (8). Such recalcitrant substrates likely constitute the primary carbon source in deeper sediment layers where labile inputs from the water column have been depleted.

In addition to these dominant taxa, we also observed a large fraction of CAZyme transcription originating from contigs not assigned to MAGs. Although these unbinned CAZymes typically exhibited lower per-gene expression values, their cumulative transcription signal indicates that carbohydrate degradation is likely distributed across a broader diversity of community members (**Figure S9**). However, the identified taxa likely are the primary functional drivers of carbon turnover, given their disproportionately high per-gene expression levels.

Together, these results suggest that the degradation of complex carbohydrates is a key process in deeper sediment layers, driven primarily by taxa within the Bacteroidota and Planctomycetota that harbour extensive CAZyme repertoires. The pronounced upregulation of carbohydrate-degrading enzymes at depths where methanogens are most transcriptionally active raises the possibility that these degradation products contribute directly to fueling methanogenesis, as discussed below.

### Plasmid and viral genes may contribute to enhanced carbohydrate turnover in deeper sediments

We also examined the occurrence and expression of CAZyme genes in viruses and plasmids. Both types of mobile genetic elements encoded diverse yet unevenly distributed repertoires of CAZyme genes across the sediment profile. Both plasmid- and viral-encoded GH genes showed a higher expression in deeper layers (**Figure 5**). The most transcribed viruses harboring CAZyme genes were Unclassified viruses, Unclassified Caudoviricetes, and giant viruses, including Phycodnaviridae and Mimiviridae. These viruses possessed a diverse repertoire of GH families, many of which extend beyond typical cell-wall degradation. These results indicate that carbohydrate-active functions are not confined to cellular genomes but are also carried and expressed by ecDNA.

**Figure 5.**
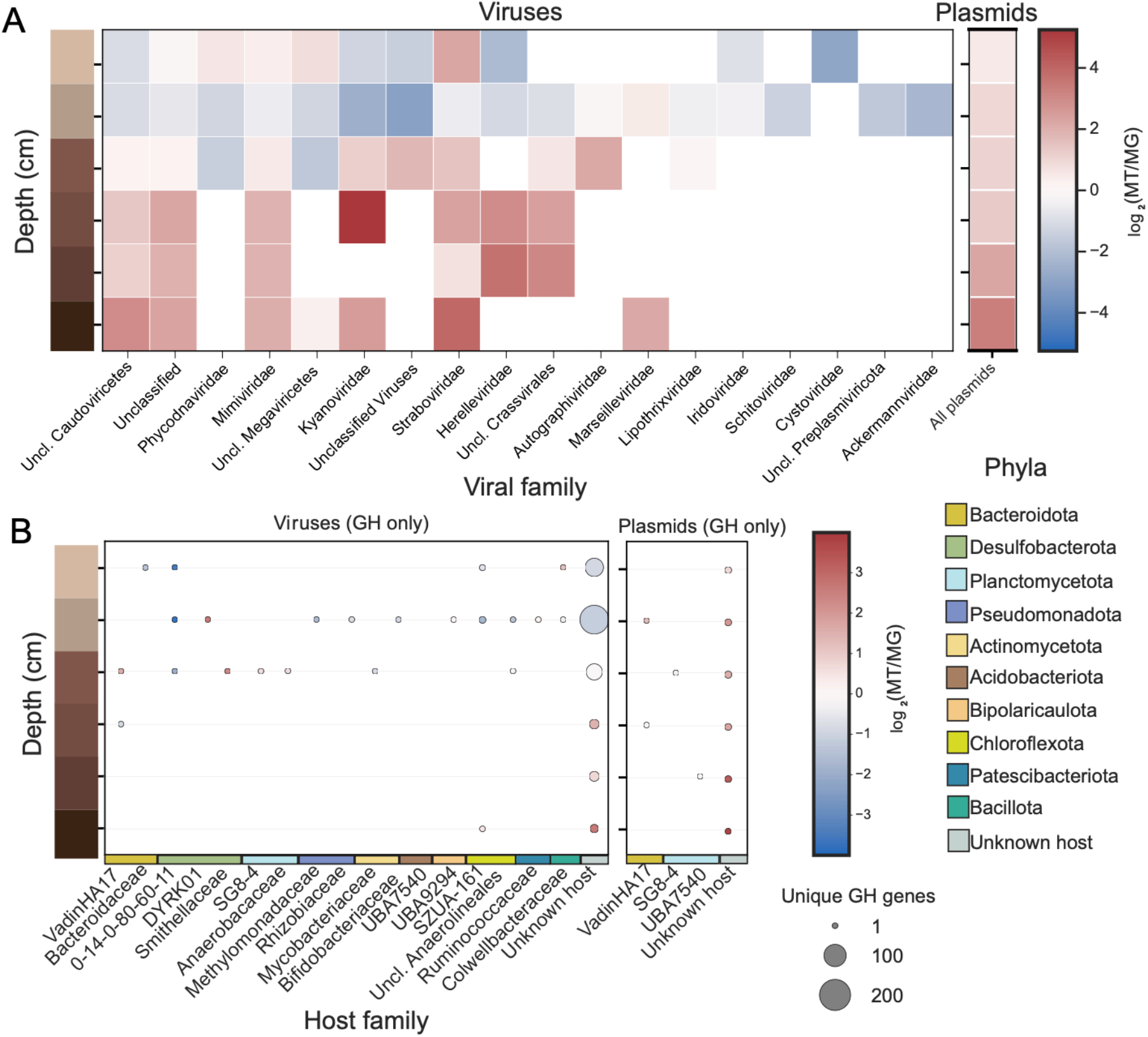
Transcriptions of carbohydrate-active gene–encoding viruses and plasmids across sediment depth. **A.** Transcriptional profiles of viruses and plasmids encoding carbohydrate-active enzymes, shown across sediment layers. **B**. Transcriptions of host-linked viruses and plasmids harboring glycoside hydrolase genes.

**Figure 6.**
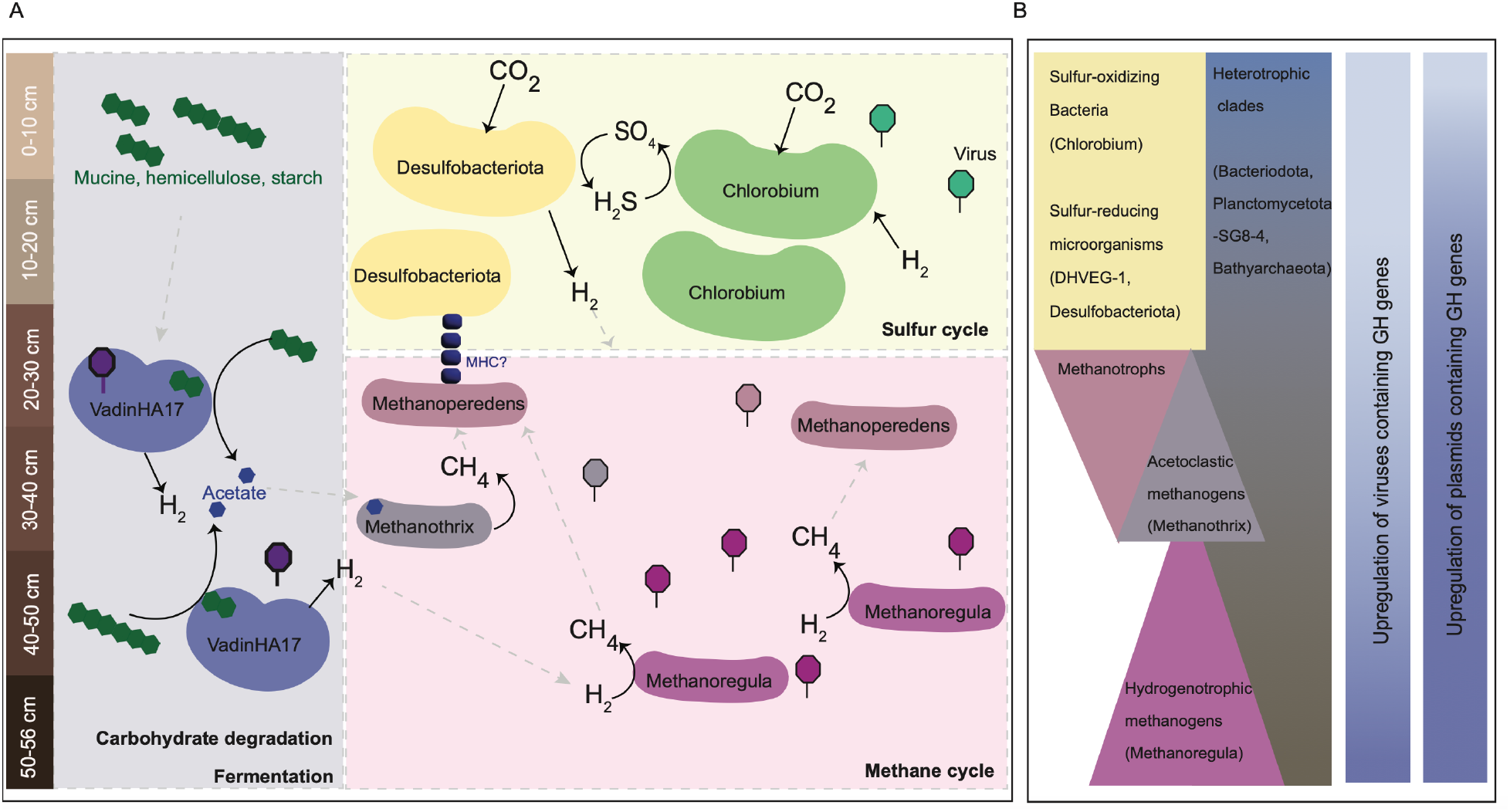
Conceptual model of microbial community organization and metabolic interactions in Lake Cadagno sediments. **A.** Cartoon representation of microbial community in the sediment, highlighting spatial structure and dominant metabolic processes. Key features include VadinHA17-mediated polysaccharide degradation and fermentation in deeper layers, the depth-stratified partitioning between acetoclastic (*Methanothrix*) and hydrogenotrophic (*Methanoregula*) methanogenesis, and anaerobic methane oxidation by “*Ca*. Methanoperedens”. MHC stands for multi-heme cytochromes. The duplication of several taxa including *Chlorobium*, VadinHA17 and *Methanoregula* indicates their presence across several layers of sediment. Viruses linked to key host taxa are shown across the sediment profile, with colors corresponding to their predicted host clades to illustrate host–virus associations. **B**. Major metabolic pathways along the sediment depth gradient, illustrating the flow of carbon from complex polysaccharides through fermentation products (acetate, H_2_) to methane production and oxidation. Heatmap of upregulation of GH-containing viruses and plasmids along the depth profile.

Viruses linked to hosts encoding GHs were predicted to infect SZUA-161 (*Chloroflexota*), *Colwellbacteraceae* (Patescibacteria), 0-14-0-80-60-11 (Desulfobacteriota), and VadinHA17 (Bacteroidota) (**Figure 5B**). Notably, the VadinHA17-associated virus carrying GH genes showed elevated transcription in the deeper sediment interval (30-40 cm), mirroring the depth-specific upregulation of both VadinHA17 family and their GHs (**Figure 3**). This viral genome spanned ∼35 kbp and encoded 50 predicted genes, including GH136. Members of this GH family have previously been reported in genomes of *Bifidobacterium* in the human gut as well as in deep-sea viral genomes (36,37), suggesting a potential role in mucin degradation (38). Phylogenetic analysis of GH136 sequences from CAZyDB, alongside homologs recovered from MAGs and the viral genome, suggests that MAG-derived proteins belong to one of the bacterial subclades, while the viral sequence branches near phage-associated homologs (**Figure S10**). However, several internal nodes show low bootstrap support, limiting confidence in deeper branching relationships. In addition, one small VadinHA17-linked plasmid contained a GH29 (exo-acting α-fucosidases). This gene was also upregulated in VadinHA17 genomes (**Figure 4**) and ranked among the most prevalent GH genes in the marine surface sediments (39). However, to our knowledge, its occurrence in plasmids has not been reported before.

Independent of iPHoP host predictions, taxonomic annotation of genes within viral elements combined with last common ancestor (LCA) analysis provided additional support for a second VadinHA17-associated virus (NODE_24693_length_7069_cov_2.266926) that was not recovered by iPHoP. This second virus encoded GH20, an enzyme known to hydrolyze glycosidic linkages in glycans, glycoproteins, and glycolipids (40). The GH20 family comprises three subclades (41), with MAG-derived GH20 sequences distributed across all three groups. The viral GH20 sequence falls within clade A-II and clusters closely with homologs from VadinHA17 MAGs (**Figure S11**), consistent with acquisition from the host via horizontal gene transfer. The substrate specificity of both GH136 and GH20 toward complex glycoconjugates suggests a specialized role in viral-mediated carbon processing.

Together, these observations suggest that these viral genes may contribute to enhanced carbohydrate turnover in deeper sediments.

### Complex polysaccharide degradation fuels methane production

The metabolic potential and activity of VadinHA17 MAGs suggests that polysaccharide degradation in this clade is coupled to fermentative metabolism. VadinHA17 MAGs lack a complete tricarboxylic acid (TCA) cycle, precluding complete oxidation of organic substates. Instead, these organisms appear to carry out mixed-acid fermentation. The high CAZyme gene expression, combined with actively transcribed fermentation pathways, indicates that complex polysaccharides are broken down and converted to acetate which is released in the environment. In addition to acetate production, this clade also encodes genes associated with H_2_ metabolism. VadinHA17 MAGs contain several hydrogen-evolving and hydrogen-sensing genes, among which [FeFe] hydrogenase group A13 and [FeFe] hydrogenase group C3 were the most expressed in deeper layers (**Table S9**). Hydrogen production during protein hydrolysis and amino acid degradation within this clade has been previously reported in anaerobic digesters, leading to its designation as the VadinHA17 family, *Candidatus* Aminobacteroidaceae (42). A similar ecological role may be inferred for Planctomycetota (SG8-4), which also exhibited strong GH transcription and has been shown to produce acetate and H_2_ (43,44). By producing both acetate and H_2_, these taxa likely function as primary degraders and fermenters that supply the key substrates for methanogenesis.

The methanogenic taxa *Methanothrix* (formerly *Methanosaeta*) and *Methanoregula* were among the most highly transcribed archaea in the deeper layers (30-56 cm). Their distribution and expression indicate depth-dependent niche partitioning of fermentation-derived substrates. In the 30-40 cm interval, acetoclastic methanogenesis by *Methanothrix* appears to dominate, whereas hydrogenotrophic methanogenesis by *Methanoregula* becomes the primary pathway in deeper layers. This partitioning is ecologically consistent with the known substrate affinities of these genera. In sediments of freshwater lakes, the H_2_ and acetate concentrations in methanogenic zones are generally low, around 10 nM for H_2_ (45) and 5-60 μM for acetate (46–49). By contrast, H_2_ concentration in the rumen can be up to three orders of magnitude higher than those observed in freshwater sediments (45,50). Acetate concentrations in lake sediments are close to the minimum threshold required for uptake by *Methanothrix* (7-80 µm acetate) (51), indicating that methane production rates are often constrained by substrate availability (52). This constraint helps explain the ecological importance of the heterotrophs and fermenters that co-occur with methanogens in these layers. By continuously regenerating acetate and H_2_ from recalcitrant polysaccharides, VadinHA17 and other fermenters may sustain methanogenesis under otherwise substrate-limited conditions.

Depth-stratified partitioning between acetoclastic and hydrogenotrophic methanogenesis has been reported across diverse freshwater and meromictic lake sediments including Lake Chaohu (29), Lake Vrana (30), Lake Geneva (53), and Lake Onego (54), Siberian lakes (55), subarctic Lake Svetloa (56), and Lake Pavin (57), where *Methanoregula* and *Methanothrix* are major lineages. The co-occurrence of active VadinHA17 (genus LD21) MAGs from a wide range of methane-rich environments including anaerobic digesters, wetlands, peatlands, methanogenic bioreactors, and lake sediments (**Figure S12**), further supports the interpretation that this clade plays a conserved role in fueling methanogenesis across diverse anoxic ecosystems.

Although VadinHA17 and methanogen transcription were not directly connected in our co-occurrence network analysis (**Figure S13**), this does not preclude metabolic coupling. Correlation-based network analyses detect coordinated fluctuations across samples, but metabolic interactions that operate at steady state may produce stable ratios, rendering them invisible. The complementary evidence from metabolic reconstruction (incomplete TCA cycle, active fermentation and hydrogenase expression), substrate specificity (VadinHA17 produces precisely the substrates that the dominant methanogens consume), and spatial co-localization of both clades and their viruses (**Figure S14**), collectively support a model of coupled carbon flow from polysaccharide degradation through fermentation to methanogenesis.

Potential for methane oxidation within the sediment was restricted to the *Candidatus* Methanoperedens (formerly ANME-2d) representing the only anaerobic methanotrophic archaeon detected in the dataset. Anaerobic oxidation of methane by *Ca*. Methanoperedens has been previously reported in the Lake Cadagno sediments (58). Recent work has also postulated that *Ca*. Methanoperedens potentially interacts with a specific sulfate-reducing bacterium, *Desulfobacteriota* QYQD01 (7), mirroring classical sulfate-dependent AOM common in marine sediments (59). While we did not find a co-occurrence, there was a coupled gene upregulation between *Ca*. Methanoperedens and sulfate-reducing QYQD01 (**Figure S15)**, rendering this an interesting aspect to be further investigated. The vertical overlap between methane production and methane oxidation suggests that *Ca*. Methanoperedens acts as a biological methane filter, consuming methane produced at depth.

## Conclusion

These results provide novel insights into a complex microbial community structured along the sediment profile. The sulfur cycle is primarily mediated by non-phototrophic sulfur-oxidizing *Chlorobia* and sulfate-reducing Desulfobacteriota. Complex polysaccharide degradation and fermentation are driven by Bacteroidota and Planctomycetota clades, which express and upregulate a diverse set of GH enzymes with depth, in order to break down increasingly recalcitrant carbon sources. Fermentation products fuel sequential acetoclastic then hydrogenotrophic methanogenesis by *Methanothrix* and *Methanoregula*, with upward-diffusing methane subsequently consumed by *Ca*. Methanoperedens. EcDNA further expands carbon degradation potential of their hosts, adding an additional layer of complexity. Overall, this study establishes a mechanistic link between early-stage carbon degradation and terminal oxidation processes throughout the sediment column.

## Data and code availability

Metagenomic and metatranscriptomic sequencing reads and metagenome-assembled genomes reported in this paper are available under ENA Project: PRJEB110274. Scripts for generating the gene catalog from metagenomic datasets and performing gene expression analysis - https://github.com/tpriest0/Profiling_metagenomes_and_mags

## Author Contributions

BD (Metagenome and Metatranscriptome analysis, Interpretation, Writing – original draft, Writing – review & editing), HZ (Conceptualization, Resources, Metagenome analysis and MAG recovery, Writing – review & editing), ES (Analysis of extrachromosomal elements, Writing – original draft, Writing – review & editing), TP (Metagenome and metatranscriptome-level analysis, Writing – review & editing), HB (Conceptualization, Resources, Writing – review & editing), KB (Resources) and MS (Conceptualization, Resources, Formal analysis, Funding acquisition, Project administration, Writing – original draft, Writing – review & editing).

## Acknowledgements

This work was supported by Swiss National Science Foundation Grant 10001203 and by the ETH Zürich. We thank Sebastian Haiss for his assistance during sample collection.

