## Supplementary figures for "Active microbial communities and their extrachromosomal elements link organic matter degradation to methane cycling in anoxic sediments"

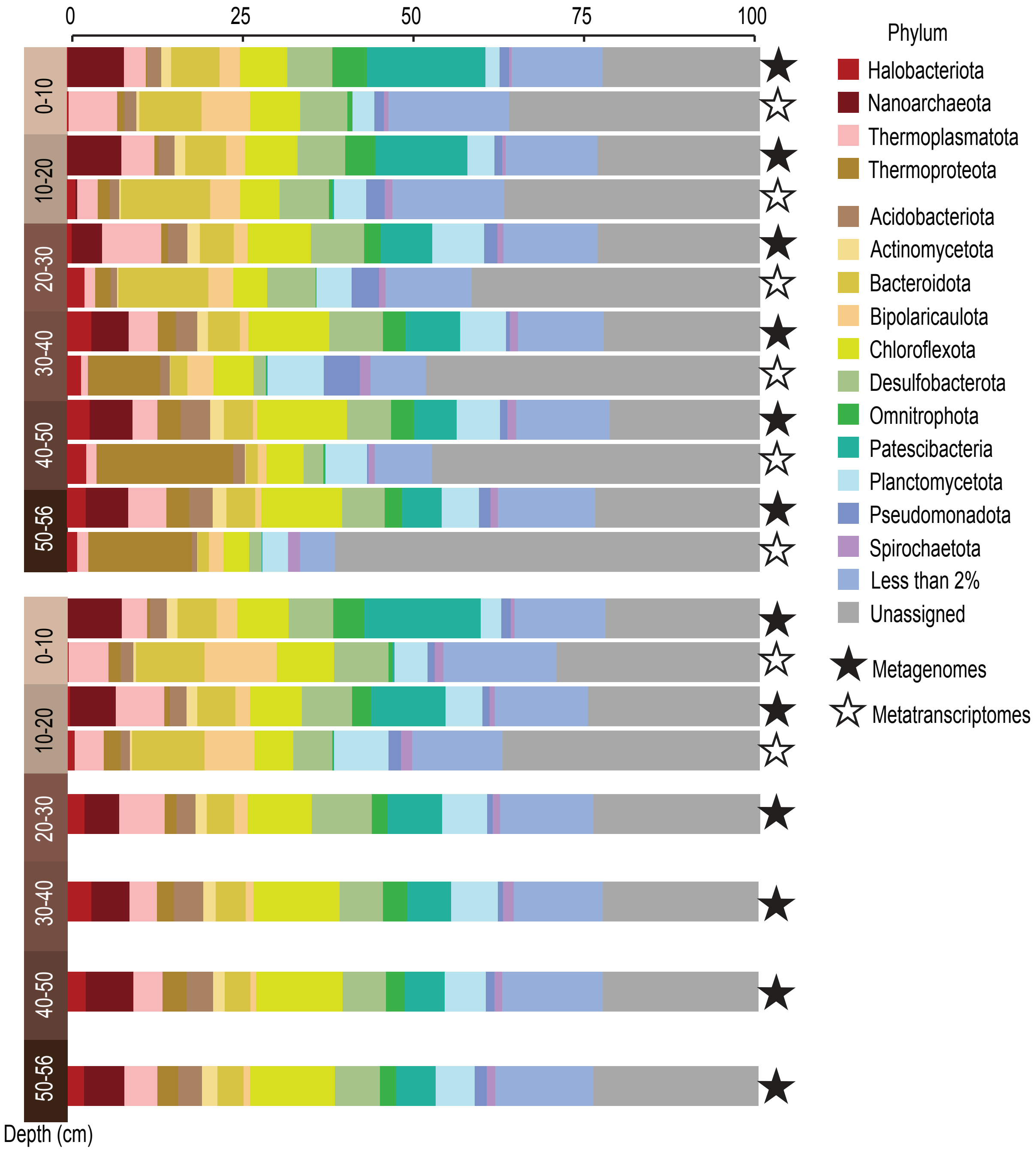


**Figure S1. Microbial community composition and transcriptional representation in lake Cadagno sediments.** Barchart showing phylum-level taxonomic composition of the total microbial community inferred from metagenomic and metatranscriptomic datasets, grouped at a 2% relative abundance threshold. Taxonomic profiles were generated using SingleM marker-gene-based profiling, and relative abundances are shown separately for metagenomes and metatranscriptomes.

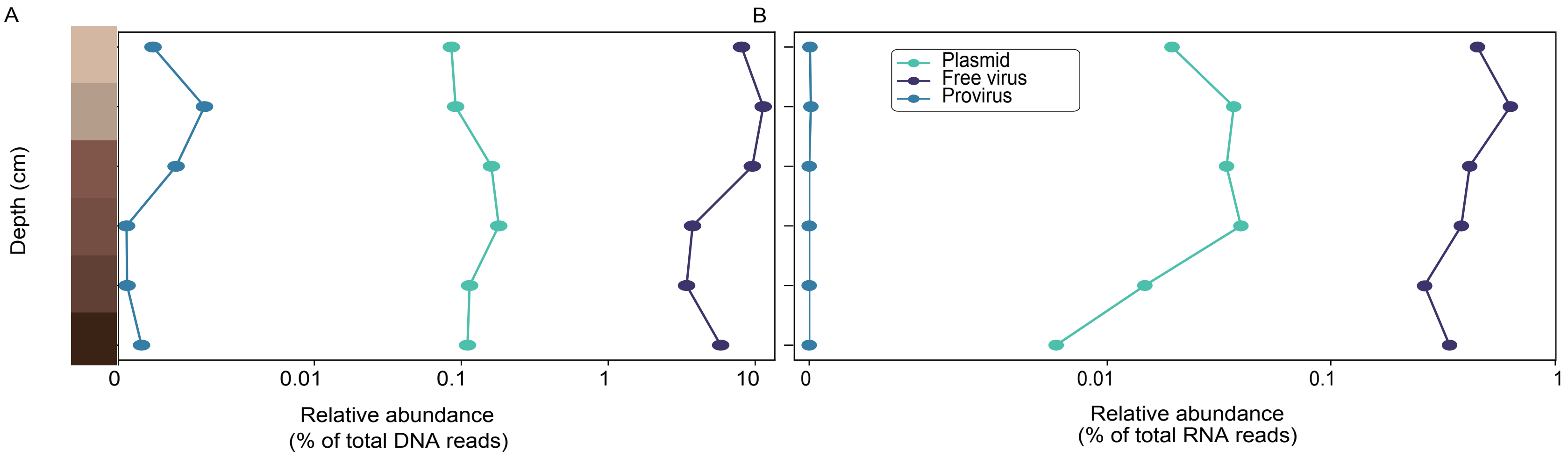


**Figure S2. Relative abundance of viruses, proviruses, and plasmids in anoxic Lake Cadagno sediments across depth.** A) Relative abundance based on metagenomic read coverage, normalized to account for sequencing depth and sample size. B) Relative abundance based on metatranscriptomic read counts, normalized to total transcript abundance per sample. Values are shown for each sediment depth interval.

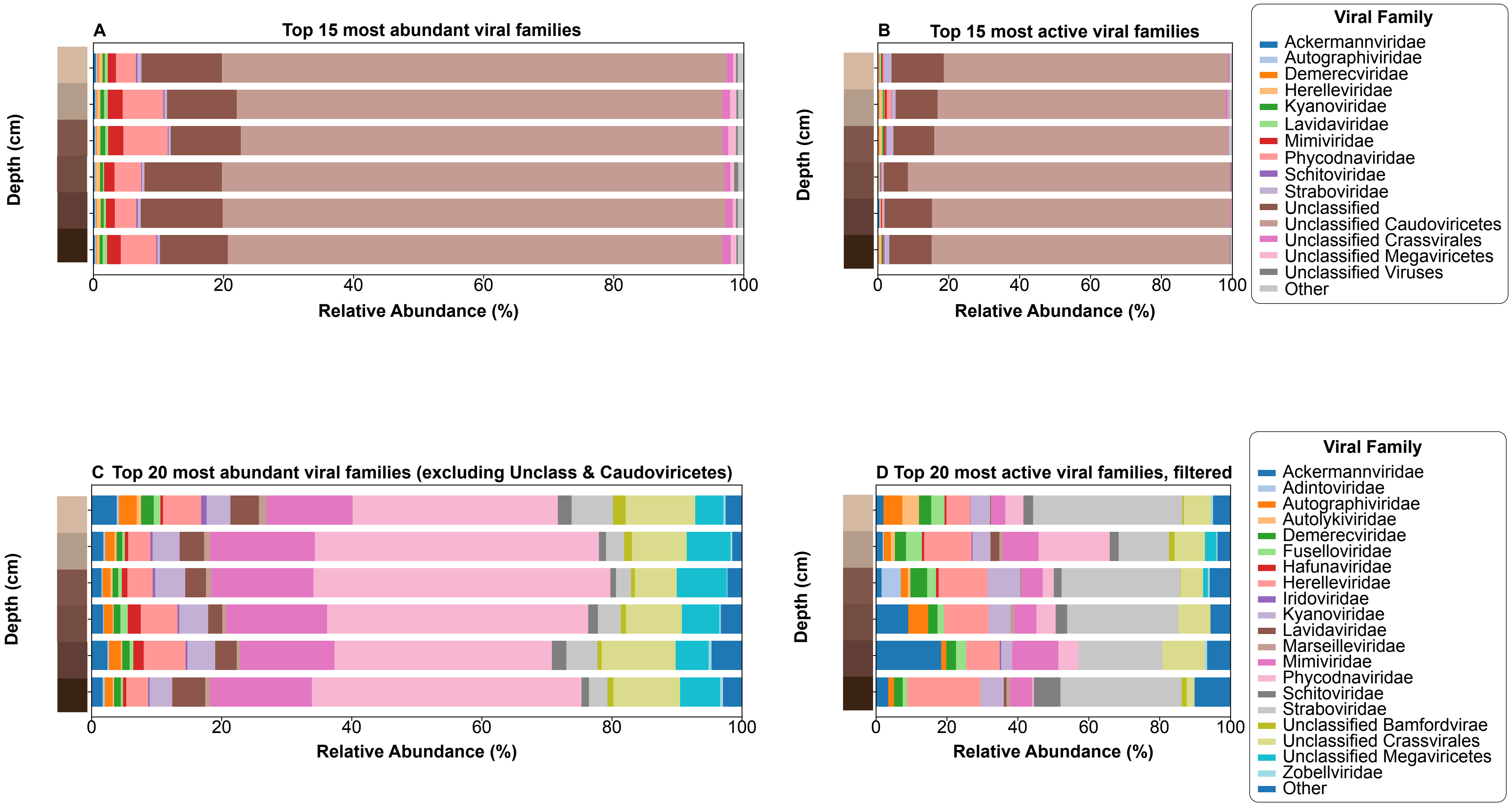


**Figure S3. Relative abundance of viruses in anoxic Lake Cadagno sediments across depth.** A) Top 15 most abundant viral families. B) Top 15 most active viral families.C) and D) Heatmaps of top 20 most abundandant and transcribed clades excluding Unclassified viruses and Unclassified Caudoviricetes.

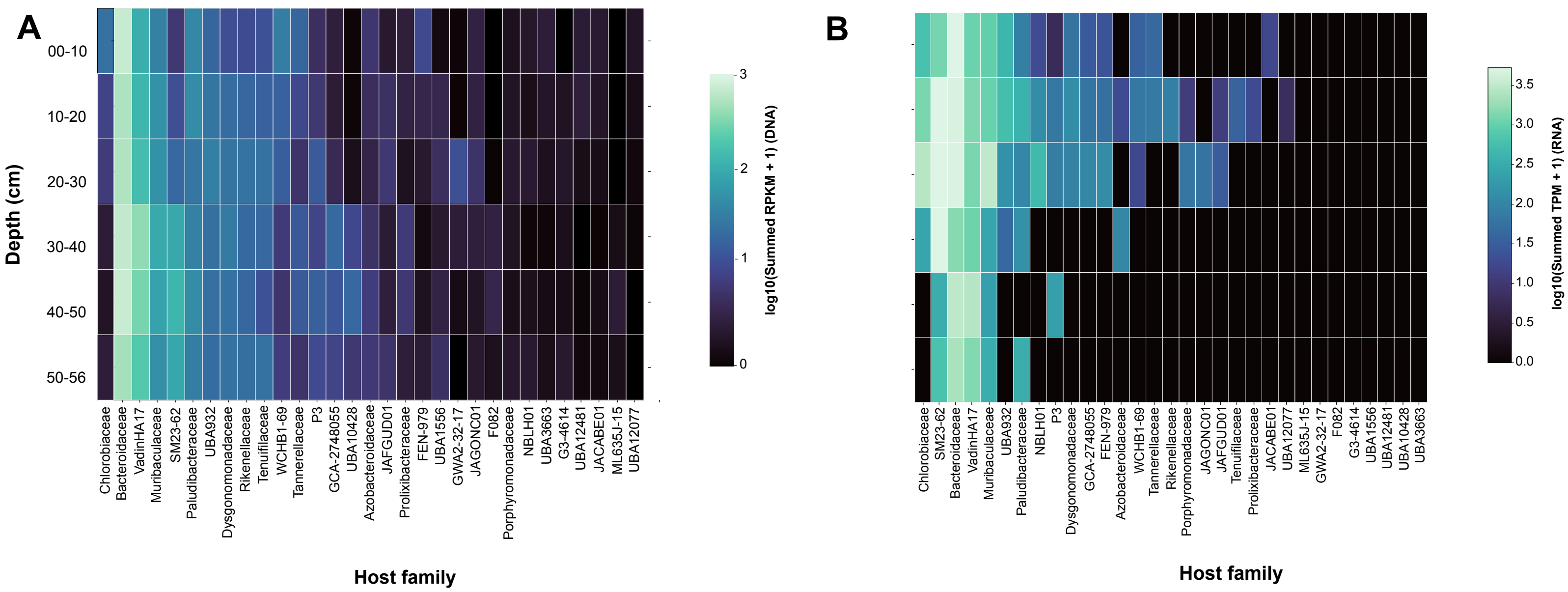


**Figure S4. Relative abundance of Bacteroidota host-linked viruses in metagenomes A) and metatranscriptomes B).**

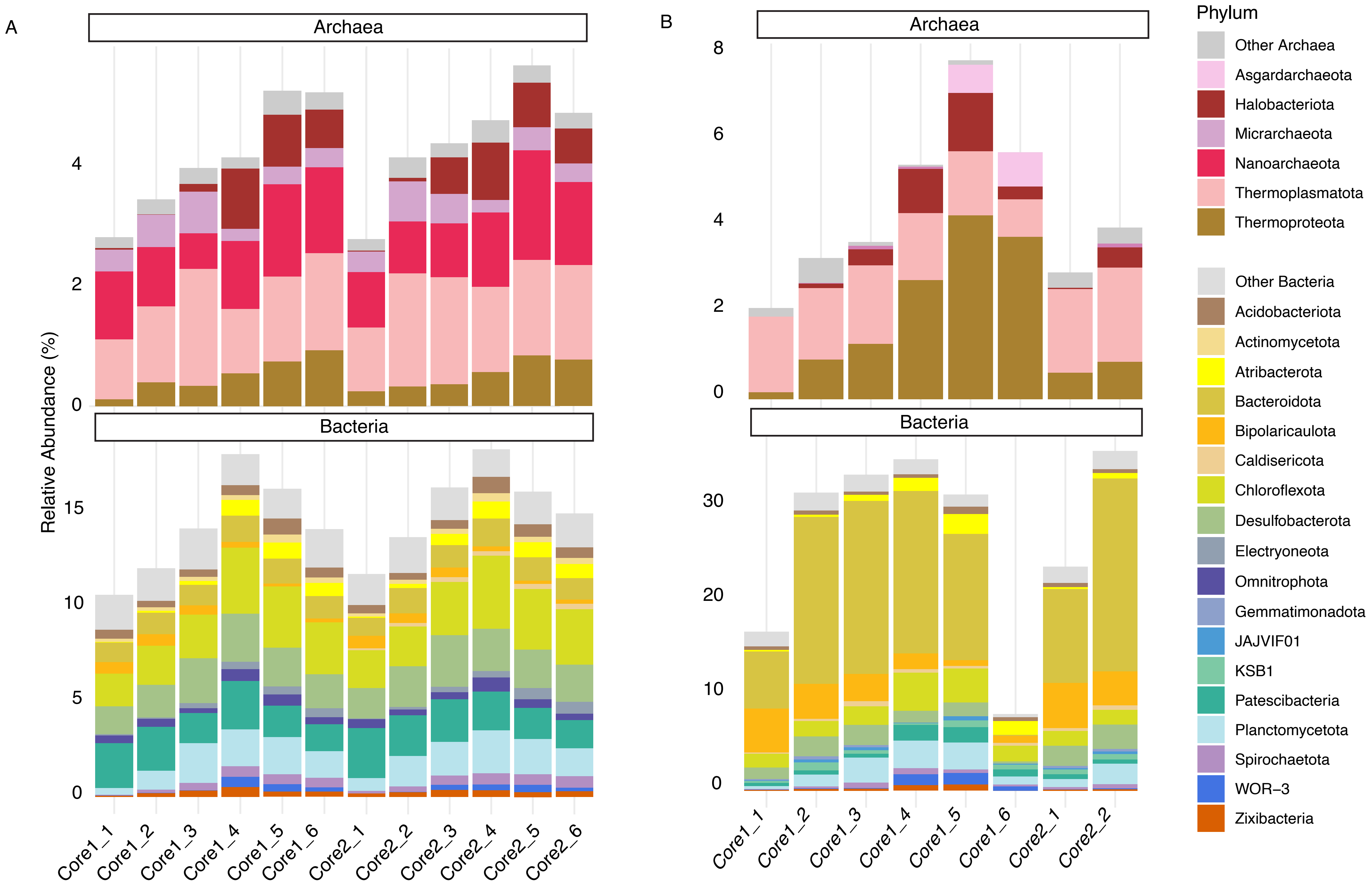


**Figure S5. Phylum-level distribution of MAG abundances across Lake Cadagno sediment depth intervals.** Analysis of abundances in metagenomes A) and metatranscriptomes B). Stacked bars show summed abundances of MAGs per depth horizon, faceted by domain (Archaea and Bacteria). The twenty most abundant phyla across the dataset are displayed individually, with lower-abundance phyla grouped as "Other."

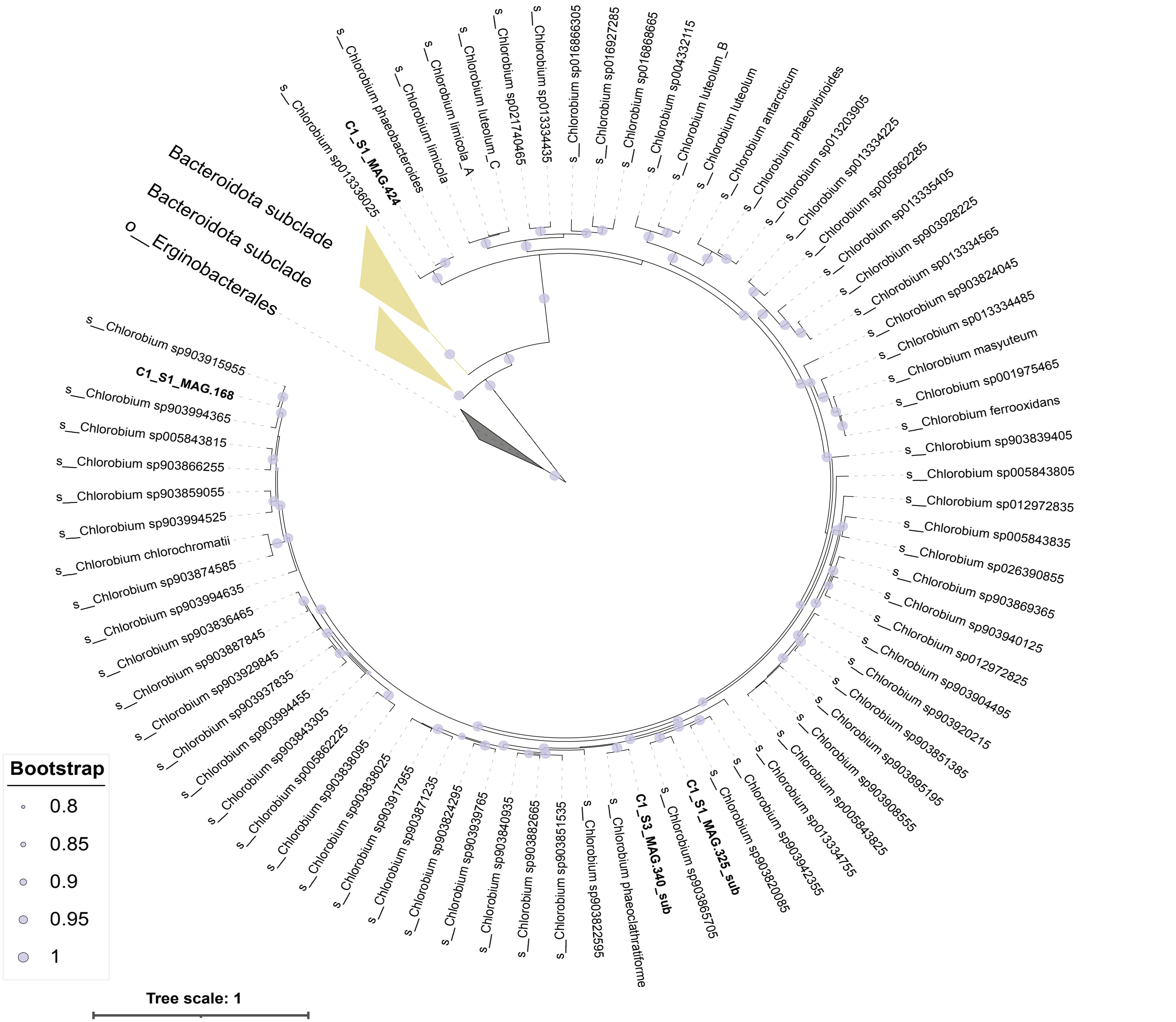


**Figure S6. Phylogenetic tree of Chlorobium (Bacteroidota) MAGs.** The tree was created using GTDB-tk with GTDB R220 database and visualized with iTOl.

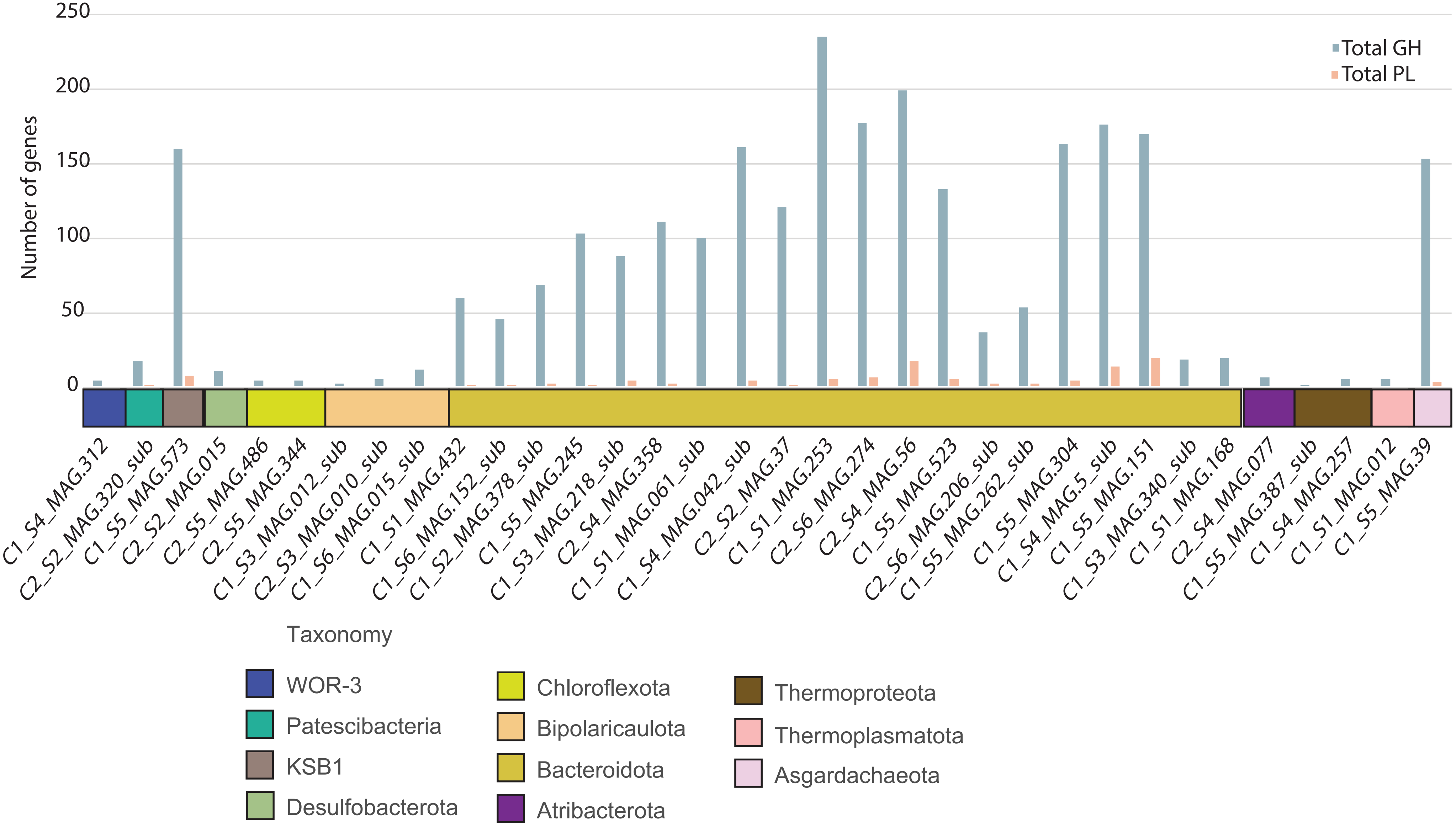


**Figure S7. Number of Glycosyl hydrolases (GH) and polysaccharide lyases (PL) in 10 most transcribed MAGs across different layers.**

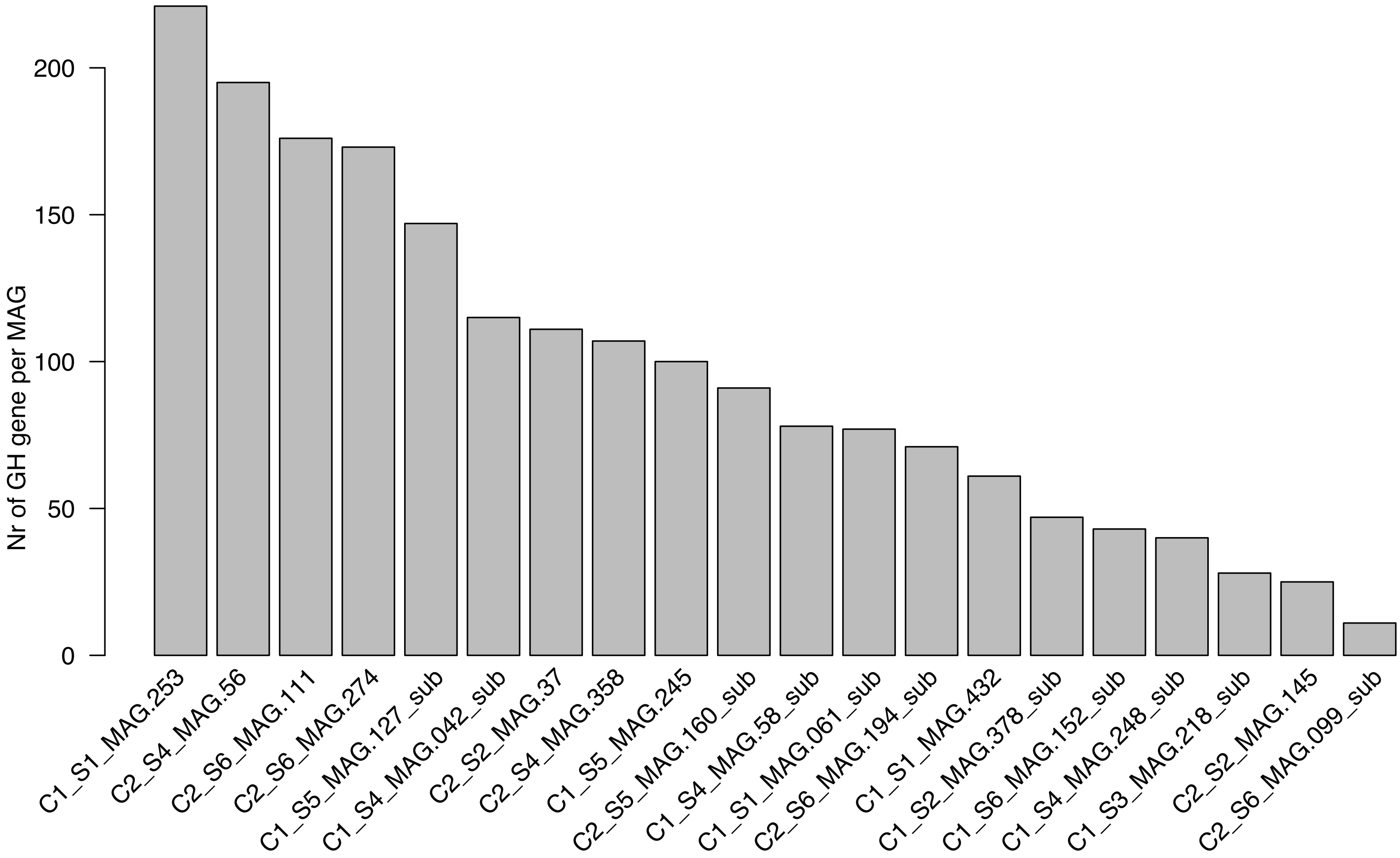


**Figure S8. Number of Glycoside Hydrolase genes annotated in each VadinHA17 (Bacteroidota) MAGs.**

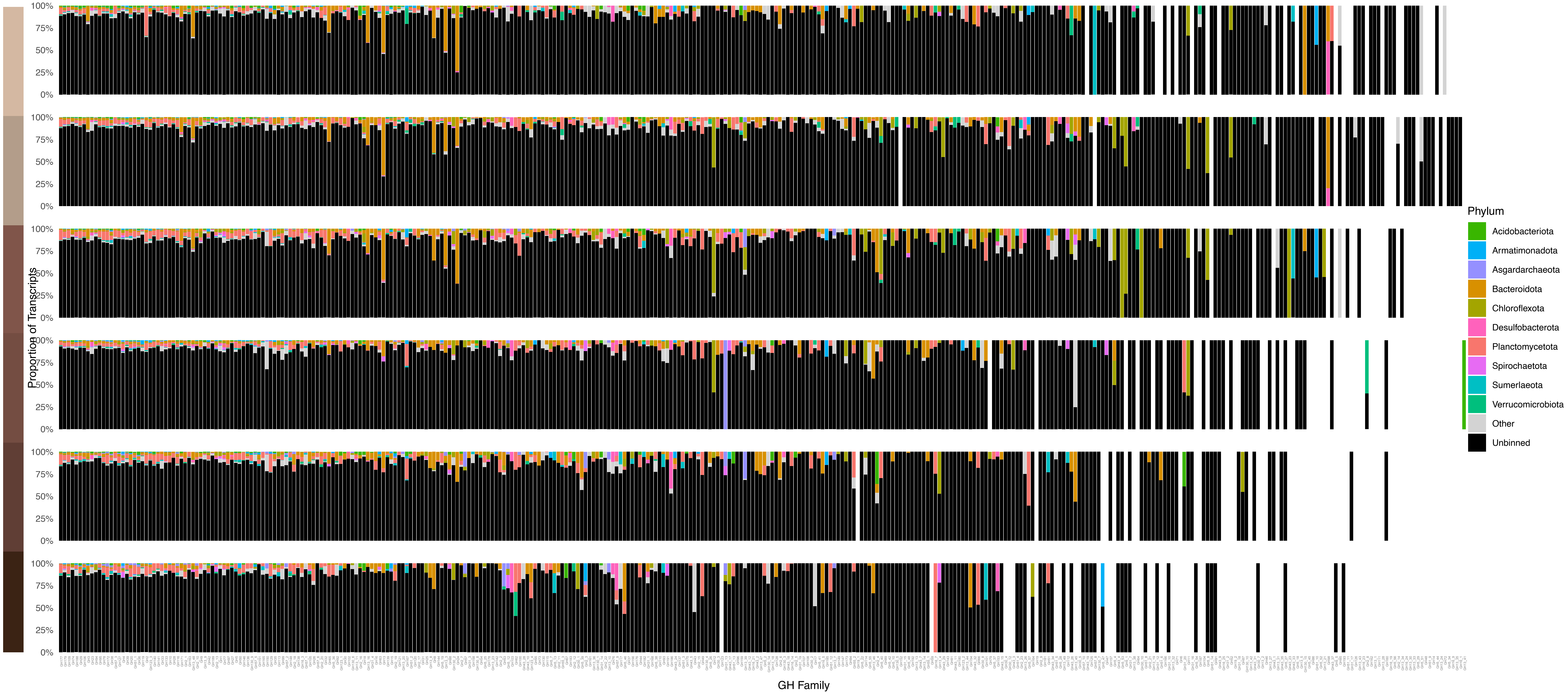


**Figure S9. Relative taxonomic contribution to the total transcript pool of Glycoside Hydrolase (GH) families across sediment layers.**

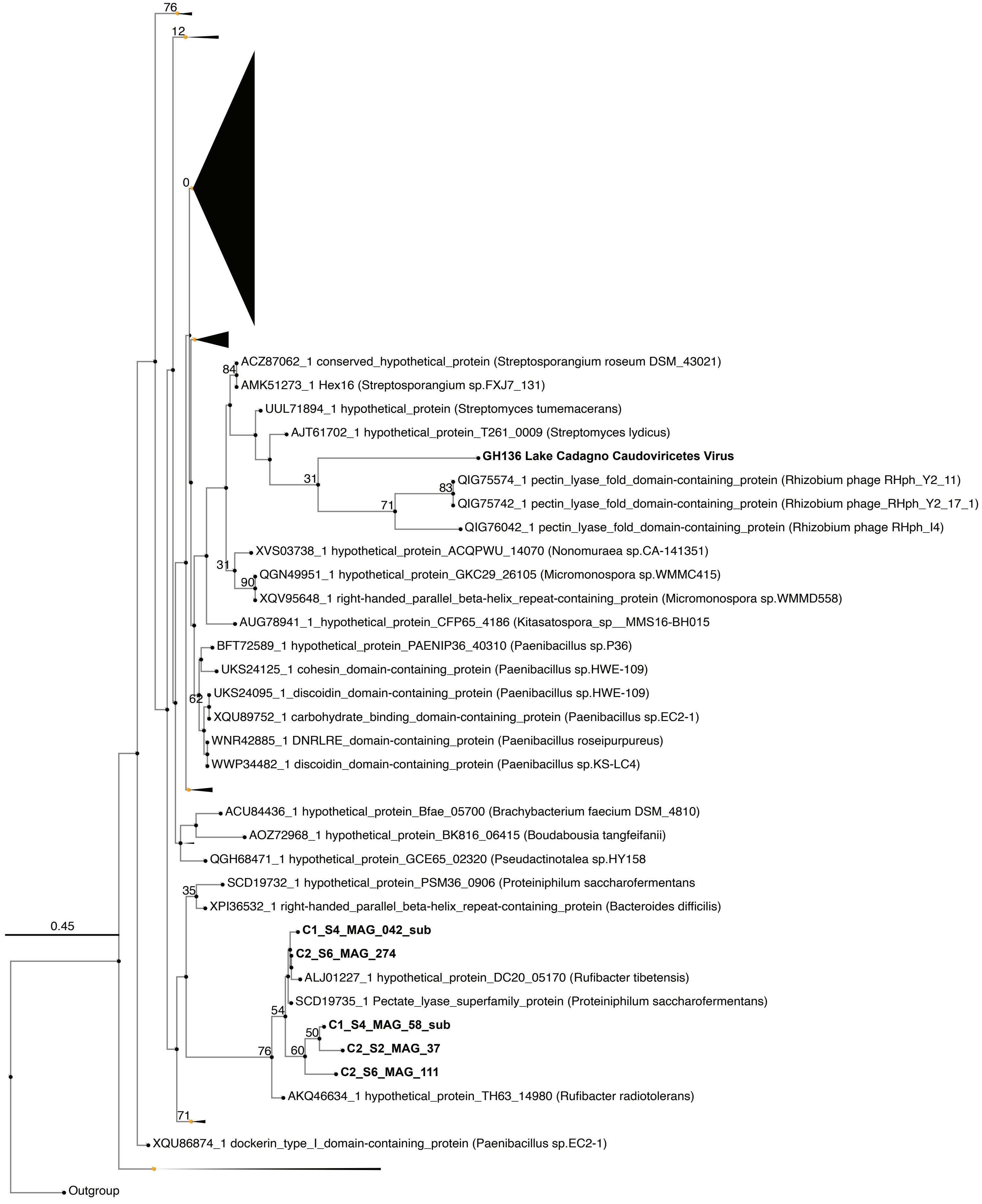


**Figure S10. Phylogenetic tree of GH136 genes, including 1 viral gene, 5 VadinHA17-affiliated MAGs GH136 and 1514 published sequences.** Tree was calculated with FastTree.

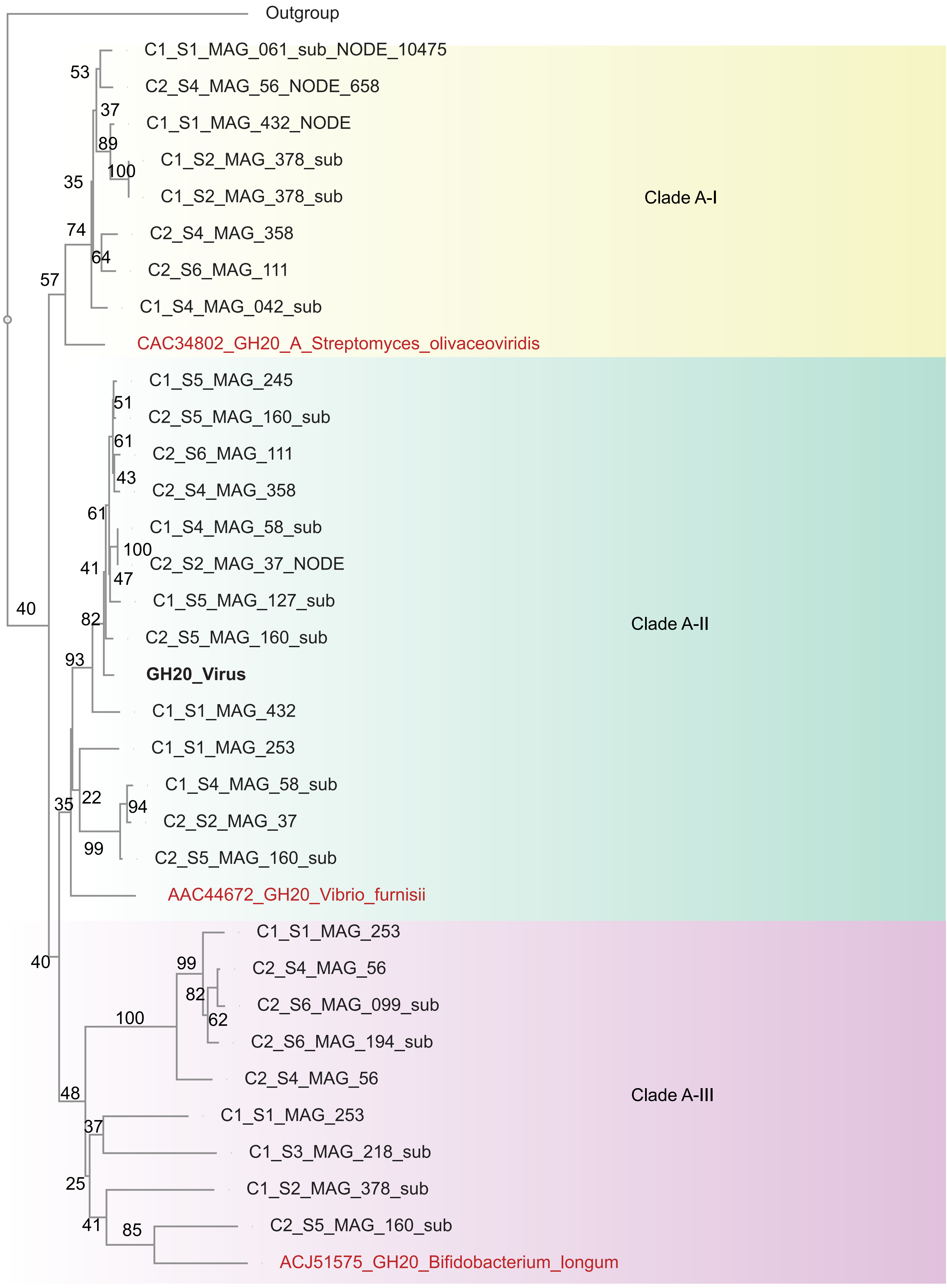


**Figure S11. Phylogenetic tree of GH20 genes including the 1 viral gene, VadinHA17-affiliated MAGs GH20 genes and published sequences.** Tree was calculated with FastTree.

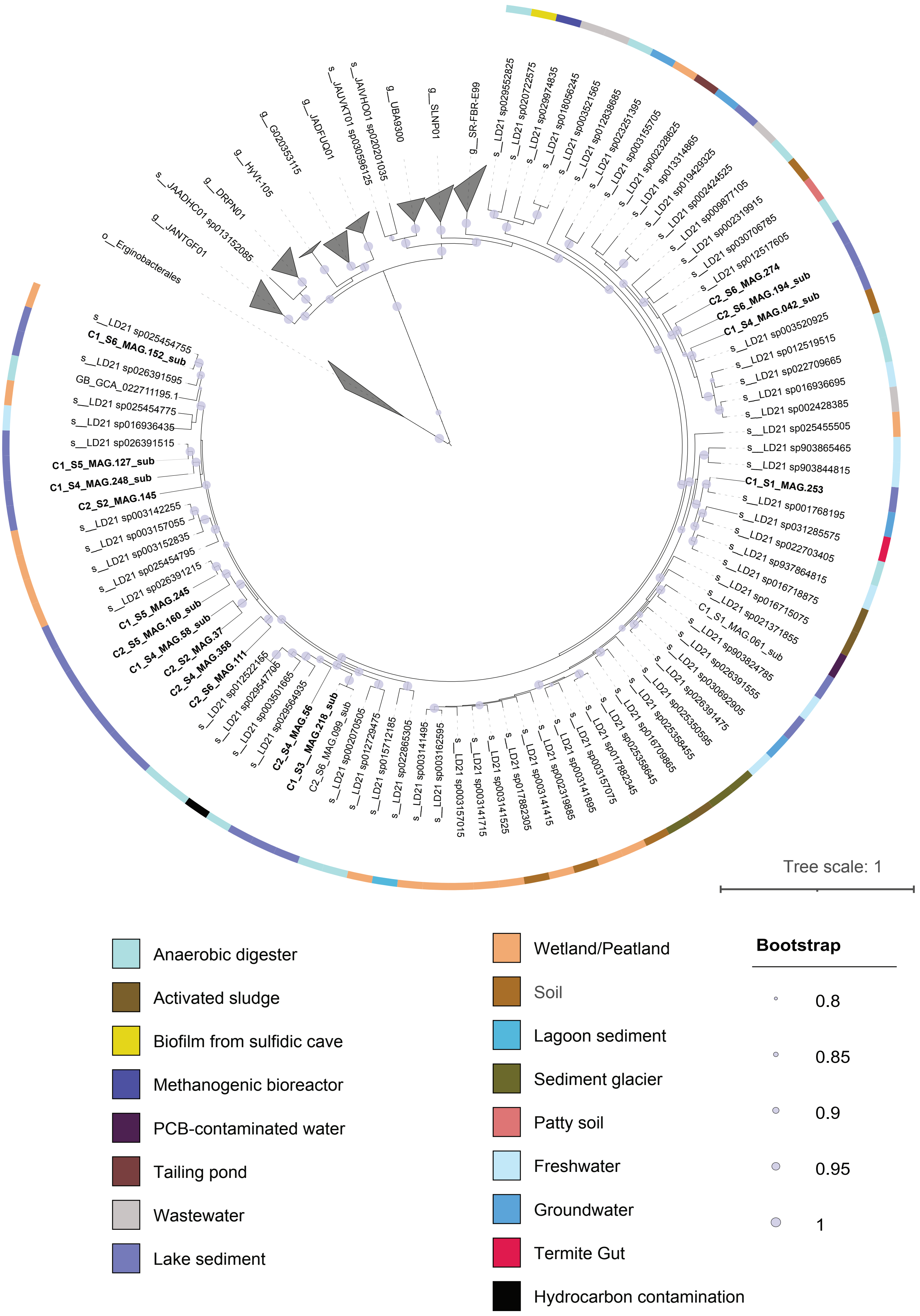


**Figure S12. Phylogenetic tree of LD21 genus (VadinHA17) MAGs.** The outer ring indicates the environmental origin of the published MAGs retrieved from the GTDB R220 database.

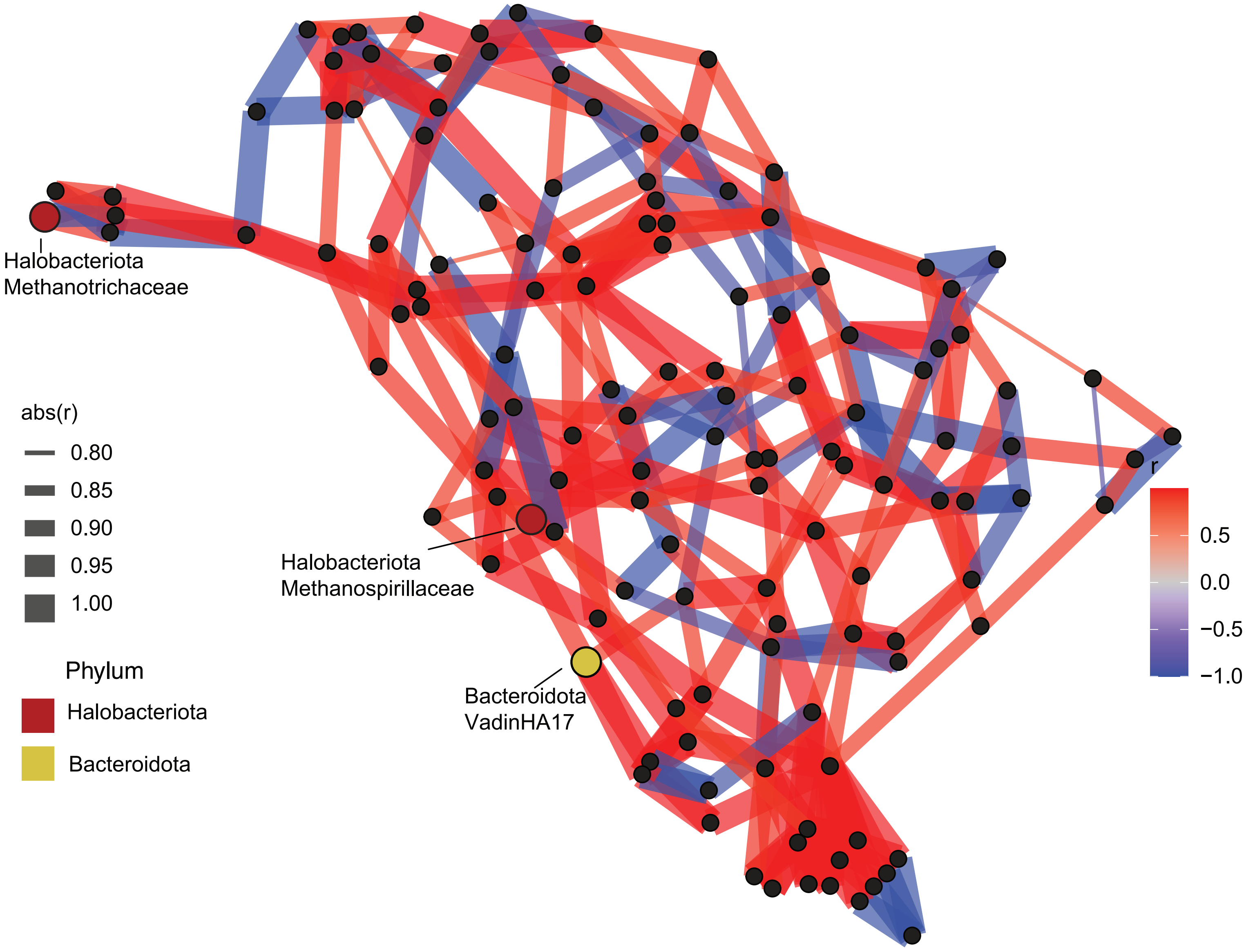


**Figure S13. Network analysis representing correlations in transcriptional activity relative to genomic abundance, calculated as $\log_2(\Sigma Mt/\Sigma MG)$.** Associations were inferred using Spearman's rank correlation, and only robust relationships ($|\rho| \geq 0.7$) were retained. For visualization, the network was pruned to the three strongest edges per node. Only taxa present in at least five samples (prevalence ≥ 5) and showing sufficient variability across samples (SD ≥ 0.3) were included.

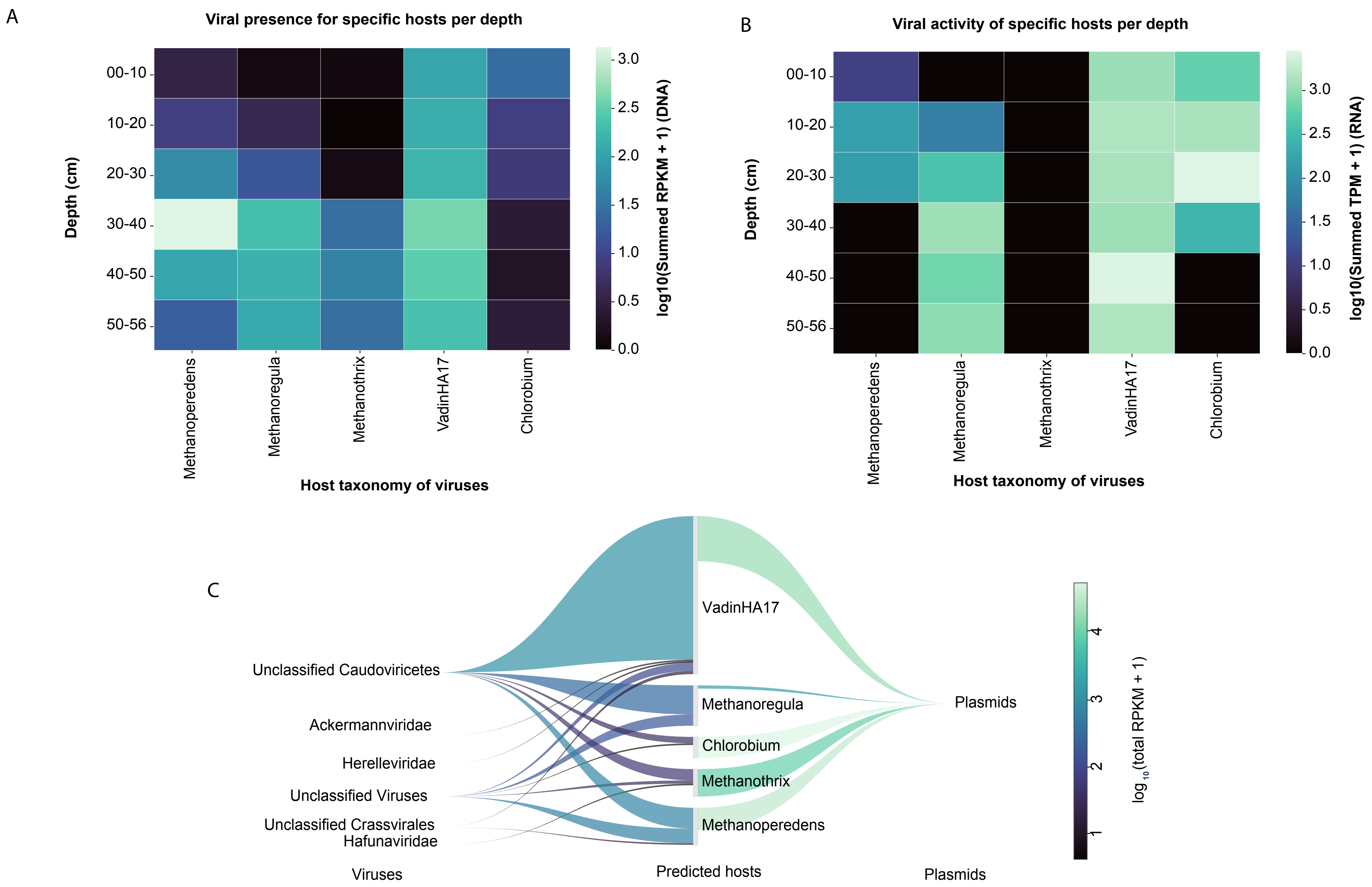


**Figure S14. Relative abundance A) and transcription B) of host-linked viruses to taxa of interest across sediment profile.** C) Sankey plot of host-linked viruses and plasmids to taxa of interest.

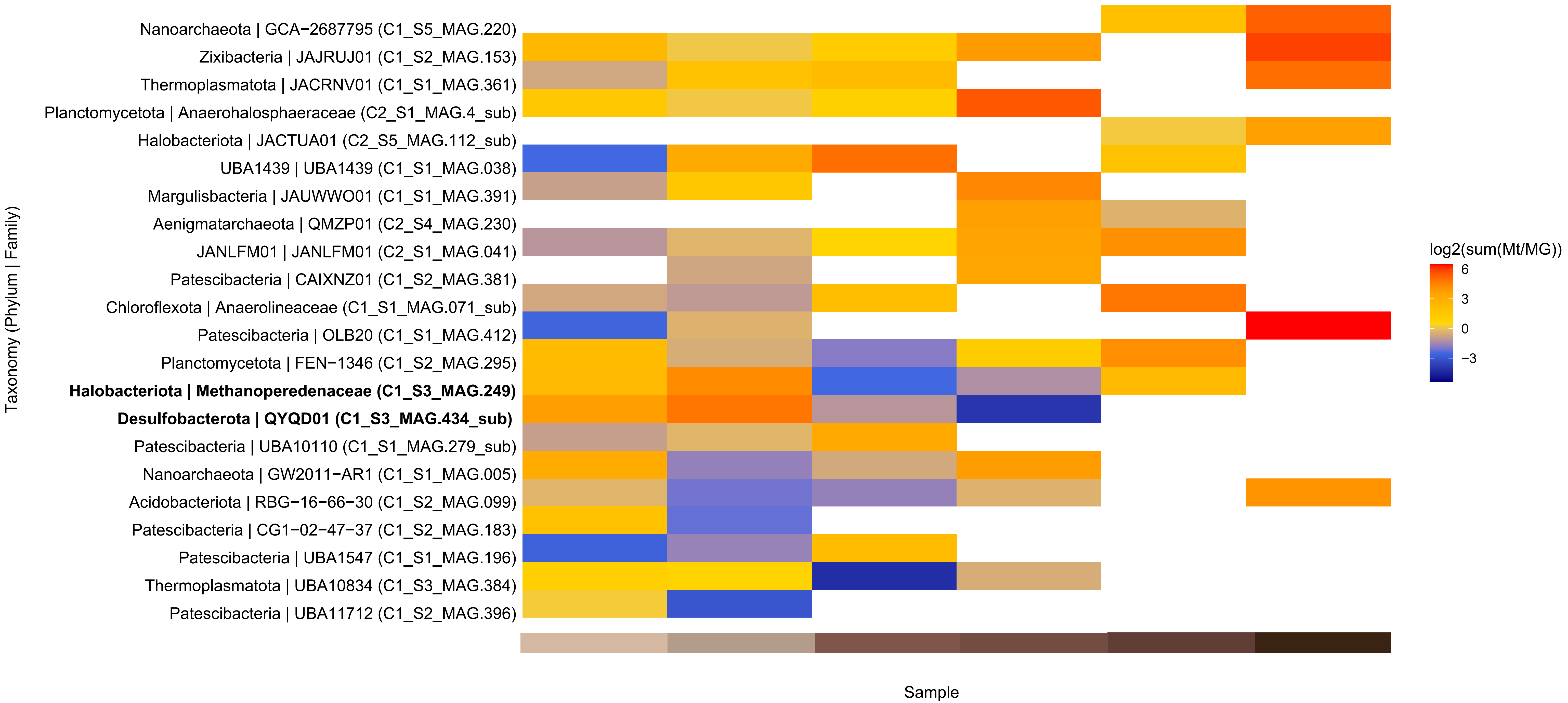


**Figure S15. Transcriptional patterns of the 25 most variable MAGs, illustrating their relative upregulation and downregulation across depths.** This analysis includes representatives from all clades and is not restricted to the top quartile of most abundant MAGs. *Ca.* Methanoperedens and *Desulfobacteriota* QYQD01 are highlighted, indicating coordinated gene upregulation at 10-20 cm depth.
