## Supplementary information for "Active microbial communities and their extrachromosomal elements link organic matter degradation to methane cycling in anoxic sediments"

**Methods**

**MAG reconstruction**

The trimmed reads were then mapped to the assemblies with BBMap v.39.01 using the parameter ambiguous=random. The resulting Sequence Alignment Map (SAM) files were converted to Binary Alignment Map (BAM) format using Samtools v.1.17 (1). BAM files were then sorted and indexed with Samtools (v.1.17). For each core, all trimmed reads from the six horizons were mapped to all six assemblies to obtain more coverage data for binning. For binning, three tools were used for each sample: MetaBAT2 v.2.17 (2), MaxBin2 v.2.2.7 (3), and CONCOCT v.1.1.0 (4). DAS_Tool v.1.1.7 (5) was used to extract the best bins from the three binning tools. Bins from all samples within each core were then pooled and dereplicated at species level (95% average nucleotide identity) using dRep v.3.4.5 (6), applying a minimum completeness of 50% and a maximum contamination of 10%, yielding medium- to high-quality metagenome-assembled genomes (MAGs) according to MiMAG standards (7) (**Table S4**).

MAGs were taxonomically classified using GTDB-Tk v.2.4.0 (8) and the GTDB database release 220 (**Table S5**). Additionally, GTDB-Tk was used for phylogenetic tree calculations of bacterial and archaeal MAGs. CoverM v.0.6.1 (9) was used to obtain relative abundances of the MAGs per sample. DRAM v.1.5.0 (10) and Metabolic v4.0 (11) was run on the final MAGs for functional annotation. Phylogenetic gene trees were calculated as described in (12).

**EcDNA identification, abundance, and activity analysis**

Viral relative abundance was estimated by mapping paired-end metagenomic reads from each depth independently to the dereplicated viral genomes using CoverM and requiring ≥95% nucleotide identity and ≥85% read alignment coverage. Coverage and abundance metrics, including mean coverage, covered fraction, relative abundance, and RPKM, were calculated. CoverM was also used to assess viral transcriptional activity by mapping metatranscriptomic reads to the same viral genome set using identical mapping thresholds and additionally reporting TPM values. Viral taxonomic identification was derived from geNomad taxonomy results. Additionally, we applied a taxonomic annotation pipeline to the viral sequences as described in Priest et al. (2023) (13). Viral host predictions were performed using iPHoP v.1.4.2 (14), with all bacterial and archaeal GTDB-Tk-classified MAGs from this study added to the iPHoP database (version Jun_2025_pub_rw).

Plasmid abundance across sediment depths was quantified by mapping quality-filtered paired-end metagenomic reads to the dereplicated plasmid reference set using CoverM. Abundance metrics including mean coverage, covered fraction, relative abundance, and RPKM were calculated using alignment thresholds of ≥95% nucleotide identity and ≥85% read alignment. To assess plasmid transcriptional activity, paired-end metatranscriptomic reads were mapped to the same dereplicated plasmid reference set using CoverM, and transcript abundance was quantified per plasmid using TPM.

**Community-level analysis: Estimation of abundance and transcription of genes, MAGs, and viral contigs**

To enable comparison of microbial community gene content across samples, we constructed a non-redundant gene catalogue. Genes were predicted from all metagenomic contigs using pyrodigal-GV v.0.3.2 (15) and clustered at 95% nucleotide identity with a minimum of 90% alignment coverage using the easy-linclust workflow in MMSeqs2 v18.8cc5c (16). Representative sequences from each cluster were retained to generate the non-redundant gene catalogue. Functional annotation of the catalogue was performed with hmmsearch v.3.4 (17) using the gathering cutoff threshold (--cut_ga) against the KEGG Orthology database (accessed March 2025) and dbCAN (version v14), with the best-scoring annotation per gene cluster selected based on bitscore and e-value.

**Co-occurrence network**

Co-occurrence networks were constructed based on the log₂-transformed ratio of metatranscriptomic to metagenomic abundance [log₂(∑MT/MG)]. Pairwise associations between taxa were quantified using Spearman’s rank correlation. Only strong correlations were retained (|ρ| ≥ 0.7). To limit network density, the three strongest edges per node were selected for downstream analyses and visualization. Taxa were further filtered to include only those detected in at least five samples (prevalence ≥ 5) and exhibiting sufficient variability across samples (standard deviation ≥ 0.3).

**Results and Discussion**

**Description of recovered MAGs**

Other abundant bacterial MAGs included Patescibacteria (n=126), Planctomycetota (n=67), Desulfobacterota (n=52), and Chloroflexota, while WOR-3 and Atribacteriota showed increased representation in deeper layers (**Figure S5**). Planctomycetota and Bipolaricaulota also contributed substantially to the transcriptomic signal. Notably, while Patescibacteria represented the most diverse phylum in terms of MAG recovery, they accounted for a relatively small proportion of total transcripts compared to other dominant taxa, in line with the observations on the metagenomic read level.

DHVEG-1 archaea are commonly inferred to be anaerobic heterotrophs, capable of degrading organic matter or fermentation products and utilizing molecular H_2_ (18,19). Their consistent high expression in the Cadagno sediments (**Table S6**) indicates an active role rather than passive persistence. In contrast, Nanoarchaeota and Microarchaeota (members of DPANN superphyla) MAGs, despite **comprising ~1% of the community**, contributed comparatively little to transcription, mirroring the pattern observed for their bacterial counterparts in the Patescibacteria (part of Candidate Phyla Radiation – CPR) (20) and in the total community analysis (**Figure 1A**). This consistent abundance-activity decoupling across both bacterial and archaeal ultrasmall-genome lineages reflects either an episymbiotic lifestyle or dormancy state, or a viral top-down regulation.

**mOTUs and SingleM discrepancies**

The two methods produced discrepant results in two aspects. First, SingleM recovered a comparatively uniform community structure across all depths, whereas mOTUs detected clear depth-dependent shifts. This discrepancy likely reflects differences in marker-gene selection and database composition. The depth gradient observed by mOTUs is more consistent with the expected decline in labile organic matter inputs with depth, which would promote shifts in heterotrophic community structure. Second, SingleM profiling recovered a striking signal from Patescibacteriota (CPR) and Nanoarchaeota (DPANN), which together comprised 5.6–16.9% and 4.3–8.0% of the community, respectively, across all samples and displayed remarkable taxonomic diversity - spanning 126 orders and 775 genera for Patescibacteriota and 366 genera for Nanoarchaeota. Despite this high genomic abundance and diversity, both groups contributed little to the metatranscriptomic signal. Whether this reflects true dormancy, episymbiotic lifestyles with low transcriptional activity, or methodological artifacts related to small genome size remains an open question and is revisited in the context of viral activity.
